# Determining cellular phylogeny and migration at high resolution in spatial genomics

**DOI:** 10.64898/2026.10.16.759405

**Authors:** Dingyao Zhang, Yunzhe Jiang, Xuezhu Yu, Donglu Bai, Jingru Tian, Archibald Enninful, Xiaolong Tian, Haikuo Li, Yiran Huo, Yuhang Chen, Nan Yao, Jiaqi Li, Zhiyuan Chu, Zhiliang Bai, Rong Fan, Jun Lu, Mark Gerstein

## Abstract

Cellular migration and tumor evolution are central to tissue homeostasis and cancer biology. Rapid advances in spatial genomics have raised the possibility of using somatic mutations to track these processes in situ in human tissues. However, frequent data dropouts in spatial genomics measurements make this challenging to implement. Here, we present StarTrak (SpaTial Analysis for lineage ReconsTRuction with somatic vAriant calling and tracKing), a framework for somatic variant calling and lineage reconstruction at single-pixel spatial resolution. StarTrak employs a neural network framework to integrate mutational evidence (from both nuclear and mitochondrial reads), spatial proximity, histological images, and transcriptomic states to identify spatially informative patterns of mutations. It performs robustly across tissue types and spatial-omics platforms (being particularly successful with spatial ATAC-seq). Applying StarTrak to tumor sections, we infer the migration of mutated tumor cells into adjacent non-malignant tissues and reconstruct distinct lineage trees associated with different spatial regions. Notably, we identify tumor regions with nearly identical transcriptomic states but distinct mutational profiles. Moreover, for skin sections, StarTrak identifies sunlight-associated mutations in non-malignant epidermis and evidence suggesting the migration of these mutant surface cells into deeper skin layers. Overall, by integrating mutation calling with spatial context, StarTrak enables the inference of spatially resolved cellular migration and cellular phylogeny.

## Introduction

A clonal population contains (nearly identical) cells descended from a common ancestral cell. Clonal evolution, characterized by the accumulation of somatic alterations and the differential expansion of descendant clones over time, is increasingly recognized as a key biological process underlying tissue homeostasis as well as tumor initiation and progression (*1*–*7*). In cancer, tumor evolution occurs within highly structured microenvironments, where factors such as tissue stiffness, immune cell infiltration, and oxygen concentration can shape the emergence and expansion of mutant cell populations (*8*–*11*). Understanding how cellular clones evolve in space and how mutated cells migrate through tissue is therefore essential for a comprehensive view of tumor progression. Although experimental lineage tracing provides a powerful means of reconstructing clonal histories in model organisms such as mice (*12, 13*), it is not readily applicable to human tissues, where lineage relationships must instead be inferred retrospectively from naturally occurring somatic alterations.

Recent advances in spatial genomics have created new opportunities to study the spatial organization of human tissues. Sequencing-based spatial profiling technologies now enable the simultaneous measurement of molecular states and spatial coordinates across thousands of microscopic spots, or pixels, within intact tissue sections, making it theoretically possible to track the evolution of mutations at high resolution in the context of the tissue microenvironment (*14, 15*). Indeed, we and others have demonstrated the feasibility of detecting somatic mutations in spatial transcriptomic data from cancer specimens using somatic mutation calls from paired bulk whole-genome sequencing (WGS) data or inferred copy number variation (CNV) (*13, 16, 17*).

However, these prior efforts were limited by major constraints. While it is feasible to detect somatic mutations in the nuclear genome, spatial genomics data are typically sparse with uneven genomic coverage, making it challenging to reliably detect somatic mutations and infer accurate phylogenetic trees at single-pixel resolution. To reduce noise, previous studies have binned pixels with similar transcriptomic states (*13*), with the underlying assumption that cells with similar gene expression must share the same mutations. Researchers have also performed pixel binning across neighboring pixels with similar gene expression for CNV analysis (*18*), with the assumption that neighboring pixels must share a uniform CNV landscape. Both assumptions are likely problematic for many tissue samples, as binning can obscure the migration of rare mutant cells into other spatial regions. Furthermore, despite the widespread use of CNV inference in such analyses, the low frequency of somatic CNV events, particularly in non-malignant tissues, limits its applicability across biological contexts (*19, 20*). For somatic mutation analysis in spatiomics data, a further challenge arises when paired bulk DNA sequencing is unavailable. Therefore, improved methods are needed to study somatic mutations at high resolution in spatiomics datasets.

To address these challenges, we developed StarTrak. StarTrak takes advantage of mitochondrial reads, which are usually discarded during spatiomics data analysis (*21, 22*), as read coverage in mitochondria tends to be much higher than that of the nuclear genome. It therefore jointly analyzes both nuclear and mitochondrial mutations at the single-pixel level. Additionally, StarTrak utilizes a multimodal autoencoder and an iterative process to refine variant profiles by integrating mutational evidence, spatial coordinates, histological features, and transcriptomic states. Existing methods for somatic mutation calling in single-cell datasets, such as Monopogen and SComatic (*21, 22*), also integrate transcriptomic information with mutational evidence, but lack the ability to utilize spatial information.

## Results

### The workflow of StarTrak

We developed StarTrak as a neural network-based multimodal embedding framework followed by an iterative refinement procedure (**Fig. 1, Fig. S1a-b**). This pipeline enables somatic mutation calling, lineage reconstruction, inference of mutation order, and characterization of mutant-cell migration patterns. We designed StarTrak for use across diverse spatial-omics technologies and specimen types. It supports both formalin-fixed, paraffin-embedded (FFPE) and fresh-frozen specimens; accommodates spatial transcriptomic and spatial ATAC-seq data; and is applicable to both spatial and single-cell datasets (**Fig. S1c-d**; see details in **Supplementary Material)**. To address the sparsity of mutational information at the individual pixel or cell level, a major challenge in spatial and single-cell datasets, StarTrak uses mitochondrial variants as an additional source of mutational information (**Fig. 1a**). Conventional spatial and single-cell analyses commonly discard mitochondrial reads, despite their substantially higher coverage than reads mapping to the nuclear genome (see details in **Supplementary Material)**.

**Figure 1.**
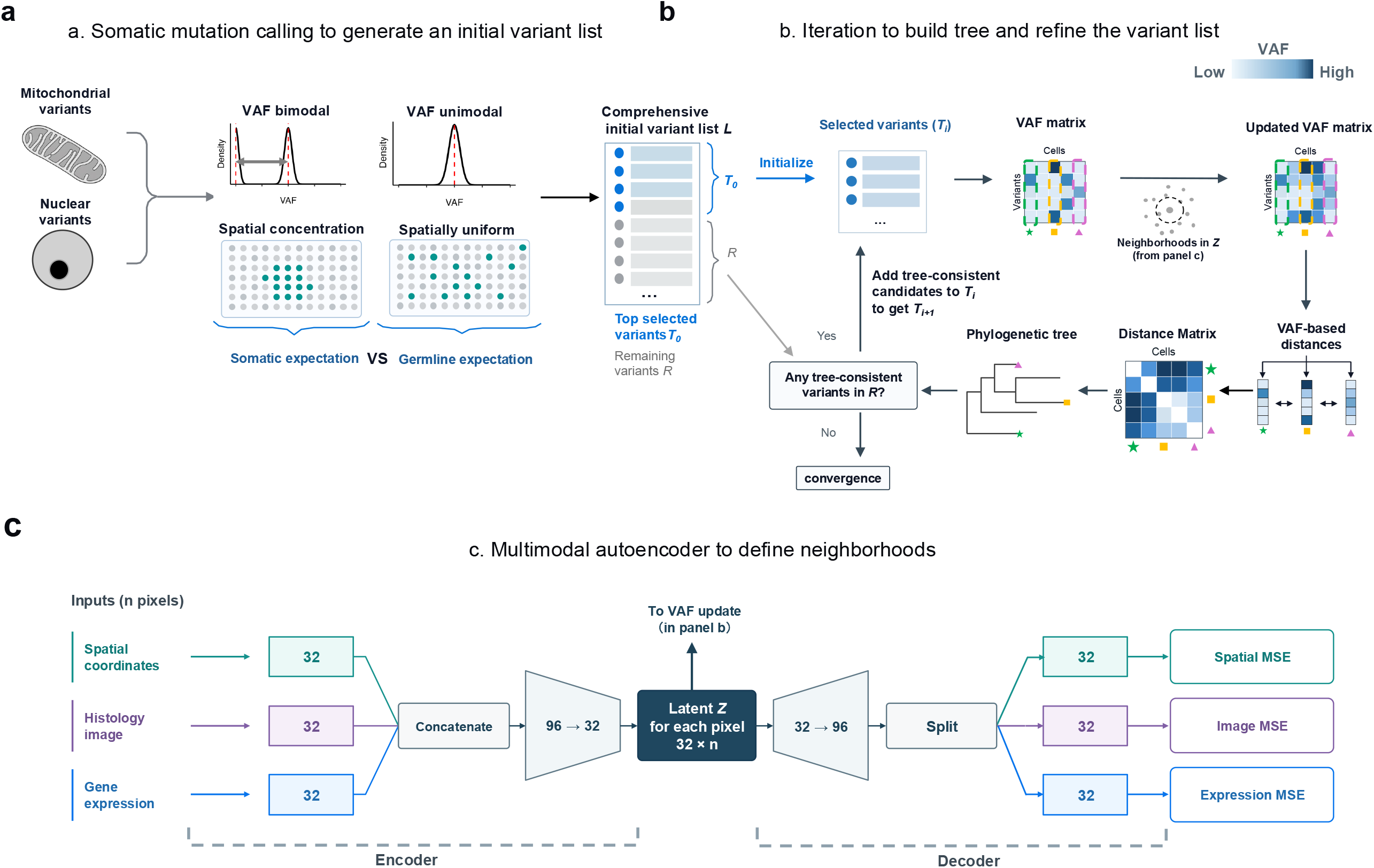
StarTrak for lineage reconstruction and somatic variant identification. **a**, StarTrak uses spatial concentration and variant allele frequency (VAF) separation to identify candidate somatic variants, generating an initial variant list for downstream analysis. **b**, RNA variant profiles and the multimodal embedding are jointly modeled for somatic mutation calling and lineage reconstruction. **c**, Multimodal inputs (including gene expression, spatial location, and histological image) are integrated through an autoencoder to produce spot-level representations. MSE: mean squared error.

StarTrak comprises three main computational components: (a) somatic mutation calling to generate a comprehensive initial variant list; (b) iterative tree building and variant list refinement; and (c) a multimodal autoencoder to define neighborhoods.

<u>(Component a) Mutation Calling</u>. The first component addresses a key challenge: most RNA variants detected by high-throughput sequencing, whether of nuclear or mitochondrial origin, are germline variants or technical artifacts, with only a small fraction representing somatic mutations (*21, 22*). To enrich for informative candidates, StarTrak prioritizes potentially somatic variants using two complementary signals, without requiring an inferred phylogeny **(Fig. 1a, Fig. S1a)**. The first captures heterogeneity in variant allele fractions (VAFs) across pixels. For germline variants, alternative-allele (ALT) reads are expected to arise broadly across covered pixels, producing VAF distributions (i.e., “read-stack” depths) that are largely consistent with binomial sampling. By contrast, somatic variants are restricted to subsets of cells or clones and are therefore expected to deviate from the binomial expectation, likely giving rise to a bimodal VAF distribution. In such a distribution, the peak near zero corresponds to mutation-negative pixels, whereas the peak at higher VAFs corresponds to pixels containing somatic mutations. The second signal captures spatial organization. Germline variants and most technical artifacts are not expected to exhibit spatial concentration and should therefore appear spatially uniform. Somatic variants, however, are expected to display structured spatial distributions, such as localized clusters or specific tissue regions. StarTrak uses these two signals to rank variants separately, then combines highly ranked variants from either ranking into a candidate pool, removes duplicates, and excludes annotated single-nucleotide polymorphisms, RNA-editing sites, and variants that fail read-level quality control, yielding a comprehensive initial variant list, *L* (see details in **Methods**). Multimapping reads can provide valuable information, but they remain underutilized by existing methods (*21*–*23*). StarTrak also includes an optional branch that processes multimapping reads separately from uniquely mapped ones (see details in **Supplementary Material**). Applying this branch revealed complementary and spatially informative variant signals that would otherwise be missed when multimapping reads are discarded (**Fig. S2**).

(Component b) Iterative Variant Refinement and Tree Building. Next, StarTrak uses the variant list, L, in an iterative framework for phylogenetic reconstruction and somatic mutation list refinement **(Fig. 1b, Fig. S1b)**. Before the iterative procedure, StarTrak selects the initial tree-building variant set, *T*_*0*_, from *L*. By default, *T*_*0*_ is the union of up to 40 top-ranked eligible variants from the spatial separation ranking and 20 from the VAF separation ranking. At each iteration, to compensate for sparse mutational evidence at individual pixels, StarTrak propagates variant information for the current tree-building variant set, *T*_*i*_, among neighboring pixels identified in the multimodal embedding space defined in component (c). The contribution of each pixel is weighted by its read depth at the variant site, such that pixels with greater read depth exert more influence on their neighbors while being less affected by neighboring values, thereby refining VAF estimates. StarTrak then generates updated pixel-level VAFs and uses them to calculate an evidence-weighted Jaccard dissimilarity matrix, from which it constructs a phylogenetic tree using the average-linkage method. Once the average-linkage hierarchical clustering tree construction is complete, StarTrak evaluates the remaining candidate variants in *L*, which have not yet been used for tree construction, for concordance with the inferred tree. Variants supported by the tree are added to *T*_*i*_ to form *T*_*i+1*_, which is used for the next iteration. This iterative procedure continues until no additional variants are admitted (see details in **Methods**).

(Component c) Multimodal Embedding. The final component learns an embedding by integrating multiple sources of evidence, including spatial coordinates, histological images, and gene expression profiles. This provides an indirect proxy for relationships among pixels without directly using mutational information. Within an autoencoder, each available input modality is encoded into a modality-specific representation. These representations are then concatenated and projected into a shared latent space, from which the input modalities are reconstructed. After training, the autoencoder contains a multimodal embedding, Z. Each embedding dimension is standardized, and pairwise Euclidean distances between pixel-level embeddings define the pixel neighborhoods used in component (b) (**Fig. 1c**; see details in **Methods**). (In a variant of the procedure, we can also bootstrap the multimodal embedding by including the initial version of the variant list as a fourth input; see details in **Supplementary Material**).

### Evaluation of StarTrak for somatic mutation detection

To evaluate the ability of StarTrak to detect somatic mutations from spatial genomics data, we first examined a gastric mucosa-associated lymphoid tissue (MALT) lymphoma specimen and several angioimmunoblastic T cell lymphoma (AITL) samples. For these samples, we had previously generated spatial transcriptomic data (Patho-DBiT) and spatial ATAC-seq data (epi-Patho-DBiT) from the same tumors, together with WGS or whole-exome sequencing (WES) data from both tumor and normal samples from the same donor (*13, 24*). These samples allowed us to evaluate somatic mutation calls against bulk WGS/WES data as the ground truth and compare results from spatial transcriptomic and spatial ATAC-seq data.

We benchmarked mutation calling using paired WES or WGS datasets and spatial genomics datasets across platforms **(Fig. 2a)**. Compared with spatial RNA profiling, spatial ATAC-seq showed broader genome-wide coverage across sequencing depths and yielded more detected variants: spatial ATAC-seq captured 1.79%–20.1% of the genome, whereas spatial RNA sequencing (RNA-seq) covered only 0.04%–10.8%, corresponding to an overall sixfold gain (for reads with depth ≥ 5). These findings indicate that spatial ATAC-seq is better suited for somatic variant analysis. At the per-pixel level, pixels in the spatial ATAC-seq dataset contained an average of 6.5 high-confidence, bulk DNA-validated somatic variants, whereas pixels in the spatial RNA-seq dataset contained only 0.6. In addition, each high-confidence, bulk DNA-validated somatic variant was observed in an average of 36.6 pixels in spatial ATAC-seq but only 3.4 pixels in spatial RNA-seq. Consistent with this, comparison of variant composition with paired bulk DNA-seq showed that ATAC-derived variant calls were more accurate than RNA-derived calls. Among all variants with sufficient read depth, an average of 75.9% of ATAC-derived variant calls were supported by ground-truth variants from paired bulk DNA-seq, whereas the corresponding average for RNA-derived variants was 22.9% **(Fig. 2b-d, Fig. S3)**. The fraction of spatial ATAC-derived variant calls supported by paired bulk DNA-seq was likely an underestimate, as tumor heterogeneity may limit the sensitivity of detecting clonal somatic mutations in bulk samples. Together, these results show that spatial ATAC-seq offers advantages over spatial RNA-seq for spatially resolved somatic variant analysis (see details in **Supplementary Material)**.

**Figure 2.**
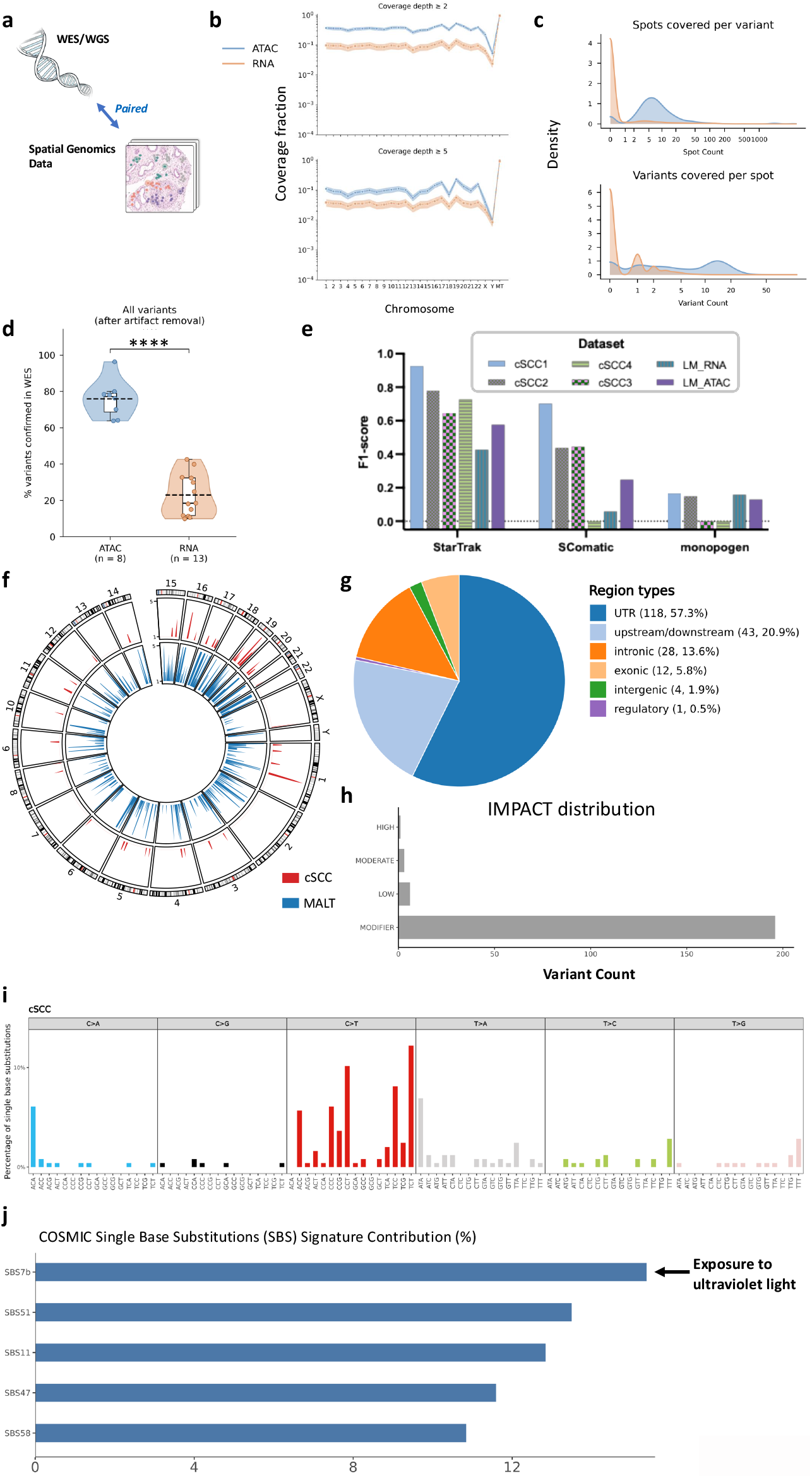
Spatial mutation analysis reveals genome-wide mutational landscapes. **a**, Overview of the benchmarking design. Whole-exome/whole-genome sequencing (WES/WGS) data were paired with spatial genomics data for mutation-analysis benchmarking. **b**, Spatial ATAC-seq and spatial transcriptomic data from the same MALT sample were compared for genomic coverage. The fraction of nucleotides covered by at least 2 (top) or 5 (bottom) reads on each chromosome is shown on a logarithmic scale. The dashed line indicates the mean, and the shaded area indicates the confidence interval. Chromosomes are shown on the horizontal axis. **c**, Density distributions of the number of spots supporting each variant (top) and the number of variants detected per spot (bottom). **d**, Comparison of variant composition in ATAC and RNA data, including support from paired bulk DNA-seq, indicating higher accuracy of ATAC-derived variant calls. ^****^ p < 0.0001. **e**, Benchmarking of StarTrak against Monopogen and SComatic across multiple datasets. StarTrak achieves higher F1 scores than the other methods. **f**, Circos plot showing the genome-wide distribution of detected mutations across chromosomes. **g**, Genomic annotation of detected mutations across different functional regions. **h**, Predicted functional impact distribution of detected mutations. **i**, Single-base substitution (SBS) mutational profile across 96 trinucleotide contexts in cSCC. **j**, Relative contributions of inferred COSMIC SBS mutational signatures.

When benchmarked against existing methods across multiple datasets, StarTrak consistently outperformed both Monopogen and SComatic **(Fig. 2e)**. StarTrak also enabled characterization of genome-wide mutational landscapes in spatial samples. Detected mutations were distributed across chromosomes but exhibited uneven variant density both across and within chromosomes (**Fig. 2f**), potentially reflecting regional variation in mutability across the genome.

Furthermore, assessment of the potential functional impact of the predicted variants revealed that most detected variants were in untranslated regions, with only one variant predicted to have a high functional impact across multiple datasets **(Fig. 2g-h)**, consistent with the expectation that most somatic mutations are passenger events. The mutational signature profiles and relative signature contributions recovered by StarTrak were also consistent with established knowledge of tumor mutagenesis **(Fig. 2i-j)**. In cutaneous squamous cell carcinoma (cSCC), the dominant mutational signature detected by StarTrak was SBS7b, consistent with its previously reported prevalence in cSCC and its association with ultraviolet (UV) radiation _(*25*)_, demonstrating the utility of StarTrak for mutational signature analysis. Collectively, these findings demonstrate that StarTrak can achieve accurate, spatially resolved somatic variant detection across diverse spatial genomics platforms.

### Evaluation of StarTrak for lineage tracing

To determine whether StarTrak can recover biologically meaningful lineage architecture *in situ*, validation in systems with known lineage structures is necessary. However, spatially resolved lineage-recording strategies remain limited. We therefore used single-cell datasets from TF-1 and A549 cells with experimentally defined clonal identities for benchmarking (*12, 26, 27*). Across the single-cell datasets, StarTrak achieved high accuracy when comparing reconstructed lineages with experimentally defined lineage relationships (**Fig. 3a-b, Fig. S4a-b**). These results indicate that StarTrak can robustly recover lineage architecture.

**Figure 3.**
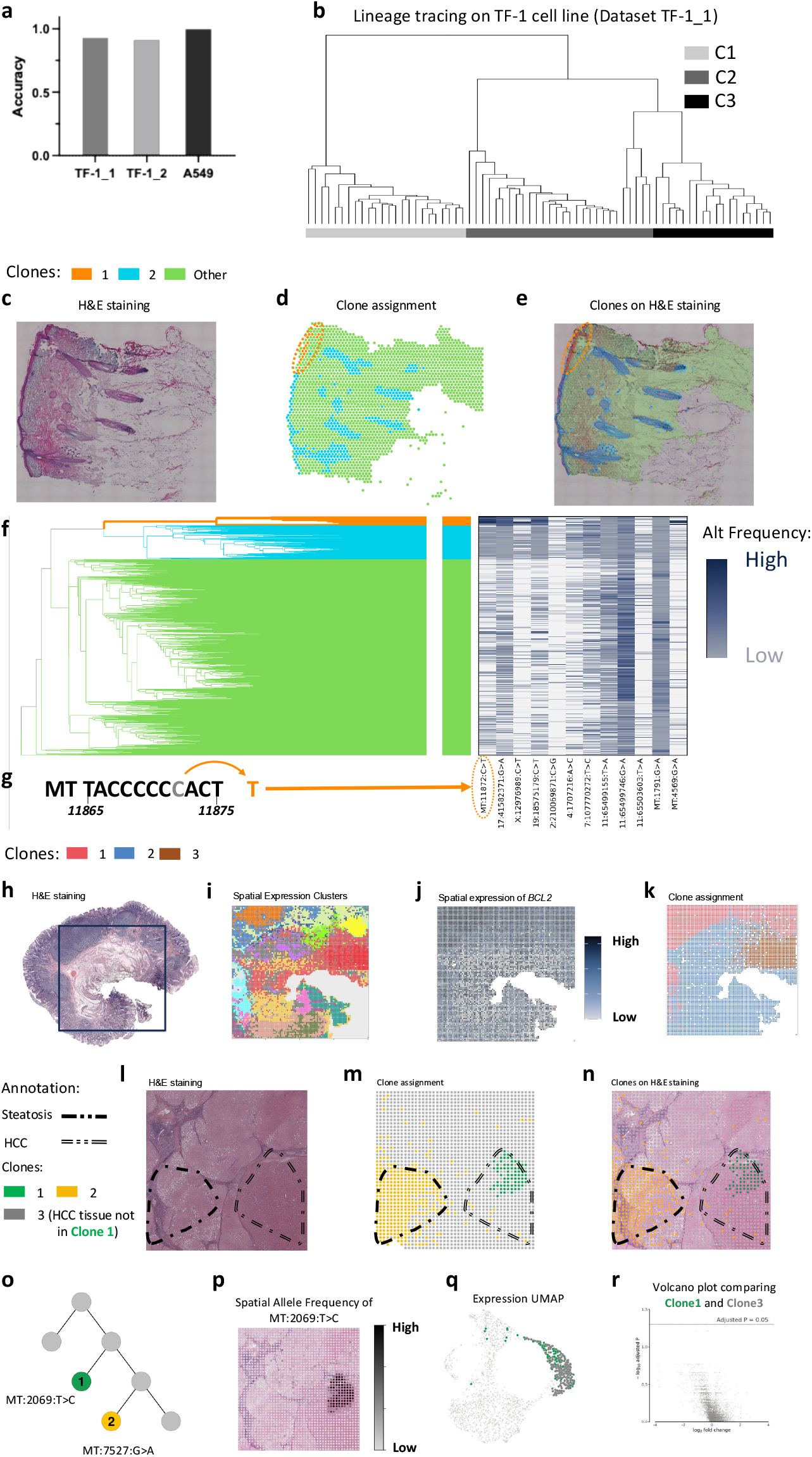
Spatial clone analysis resolves biologically meaningful clonal architecture in tumor and normal samples. **a**, StarTrak achieves high accuracy in lineage reconstructions across several datasets. **b**, StarTrak recovers ground-truth clonal structures in a TF-1 cell-line model with experimentally defined ground truth. **c-g**, Normal skin sample. **c**, Histological image of normal skin. **d**, Spatial distribution of the StarTrak-inferred clones without the histological image overlay. **e**, Spatial distribution of the clones inferred by StarTrak, overlaid on the histological image. The inferred clonal architecture successfully recapitulates the spatial organization of the skin section and identifies a distinct clone localized to the skin surface, as indicated by the circle. **f**, Phylogenetic tree and mutation heatmap demonstrate that this clone is defined by several mutations. **g**, Sequence context of MT:11872:C>T, consistent with a UV-associated C>T substitution. **h-k**, Mucosa-associated lymphoid tissue (MALT) sample. **h**, Histological image of the MALT sample. **i**, Spatial distribution of expression-derived clusters. **j**, Spatial expression pattern of *BCL2*, which may indicate the spatial extent of the B-cell lymphoma region. **k**, Spatial distribution of clones inferred by StarTrak shown alongside the histological image, showing that one clone substantially overlaps with the tumor region. **l-r**, Hepatocellular carcinoma (HCC) sample. **l**, Histological image of an HCC section. **m**, Two clonal architectures aligned with key histopathological features, including steatotic hepatocytes and HCC. **n**, Spatial distribution of the clone inferred by StarTrak with the histological image overlay. **o**, Phylogenetic relationship of those clones. **p**, Spatial distribution of the variant: MT:2069:T>C. **q**, Gene expression UMAP of the HCC sample. Green, spots assigned to the MT:2069:T>C-marked clone; dark gray, all other spots in the expression-defined cluster that contains this clone. **r**, Differential gene expression analysis comparing the clone with the remaining spots (two-sided Wilcoxon rank-sum test with Benjamini–Hochberg correction for multiple testing).

Beyond cell-line benchmarking, evaluation of StarTrak in biological specimens is necessary. Because StarTrak leverages mitochondrial mutations, which are expected to accumulate in benign tissues, we hypothesized that the method would be applicable beyond malignant contexts (*27*). Moreover, developmental relationships are often better defined in non-malignant tissues, providing a more interpretable framework for assessing lineage reconstruction. We turned to spatial RNA-seq data from a benign human skin specimen (Skin1) to evaluate StarTrak and identified multiple spatial clones across the tissue sections (**Fig. 3c–g**). Among these, one clonal lineage contained pixels corresponding to hair follicles and epidermis, distinct from the surrounding tissue compartments (**Fig. 3c-e**). This is consistent with the shared developmental origin of hair follicles and epidermis (*28*) and evidence that hair follicle-derived cells can contribute to epidermal repair following injury (*29*). Unexpectedly, StarTrak revealed another clone with a strikingly distinct spatial distribution, suggesting a potential local lineage expansion event. Notably, this clone (Skin1-Clone1) was mostly localized to a region at the skin surface and was defined by three inferred somatic variants, including one mitochondrial variant and two nuclear genomic variants. The presence of three variants in the same spatially segregated lineage argues against the possibility that this clone resulted from technical artifacts (**Fig. 3f-g**). This interpretation was further supported by the mutational spectrum of these variants, which followed the canonical spectrum of UV-induced mutagenesis, with an excess of C>T substitutions and strong enrichment at polyC sites (*30*).

### Application of StarTrak to spatial lineage reconstruction in tumors

Tumor biology is an important context for lineage analysis, as somatic mutations and clonal diversification are recurring themes. We tested StarTrak on spatial transcriptomic sections from a MALT lymphoma and a hepatocellular carcinoma (HCC) specimen (**Fig. 3h-r**). In the MALT sample, StarTrak successfully distinguished cancer-associated clones that overlapped with expression-based annotations of the tumor regions (**Fig. 3h-k**). In the HCC sample, expression-based annotations delineated regions of hepatic steatosis and carcinoma (**Fig. 3l**). StarTrak then resolved two major spatial clones that mapped to subregions containing steatotic hepatocytes and HCC cells, respectively (**Fig. 3m-n**). We next identified two mitochondrial variants, MT:2069:T>C and MT:7527:G>A (**Fig. 3o-p**), that marked distinct spatial clones. Notably, the clone marked by MT:2069:T>C exhibited a transcriptomic state highly similar to that of neighboring pixels, highlighting that phylogenetically distinct clones can share the same gene expression profiles (**Fig. 3q-r**).

Beyond clonal assignment, reconstructing the order of mutation acquisition is also important for understanding tumor evolution. By integrating phylogenetic relationships with the spatial distributions of mutations in a cSCC specimen (cSCC1), StarTrak inferred an ordered mutational trajectory (**Fig. S5a**). Together, these findings establish that StarTrak enables spatially resolved lineage reconstruction across both non-malignant and malignant tissues.

### StarTrak infers migration events in malignant and benign skin tissues

Tumor cell migration and metastasis are fundamental processes in tumor biology, and spatially resolved, high-resolution somatic mutation calls by StarTrak can provide a direct means of uncovering these processes *in situ*. We found a striking example of cell migration in cSCC1 and its biological replicate (cSCC2). A mitochondrial dinucleotide substitution, MT:12041:CC>TT, consistent with the canonical UV-associated CC>TT mutational signature, showed clear evidence of somatic origin (**Fig. 4a, Fig. S5b**). Bulk DNA sequencing supported its somatic origin, showing no evidence of the variant in matched non-malignant tissue but substantial variant read counts in the tumor sample (**Fig. 4b**). This mutation emerged as a spatially heterogeneous event that defined a distinct clone within the malignant region of the sample. Notably, many pixels harboring this mutation were also present in neighboring histologically non-neoplastic tissue (**Fig. 4c-d**), suggesting that this spatial clone migrated from the malignant region into the neighboring normal region.

**Figure 4.**
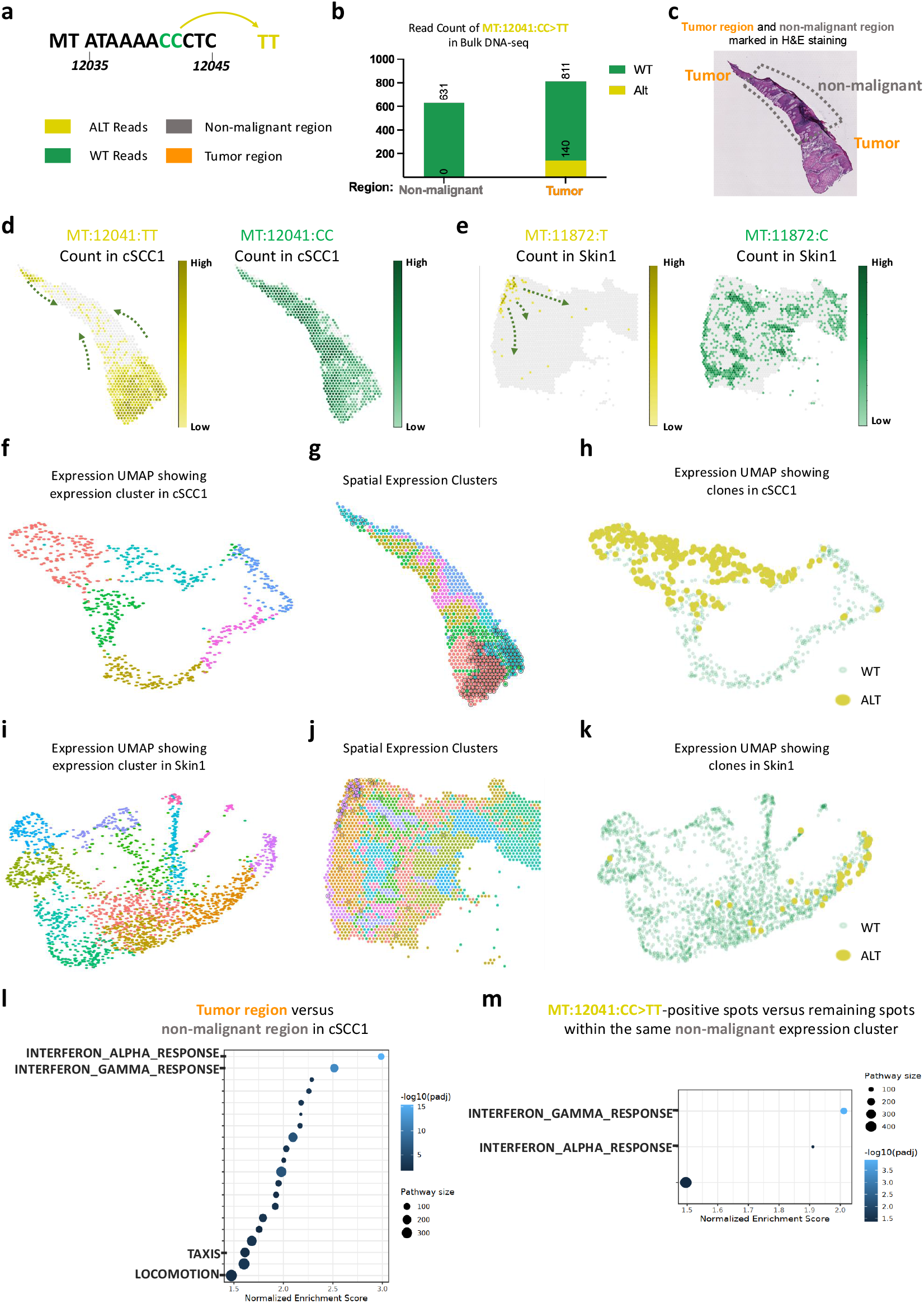
Spatial analysis reveals previously unrecognized migration of mutation-bearing cells in skin tissue. **a**, Sequence context of MT:12041:CC>TT, consistent with a UV-associated CC>TT substitution. **b**, Bulk DNA-seq validation of MT:12041:CC>TT, showing marked enrichment in tumor relative to non-malignant tissue and supporting its somatic origin. **c**, Histological image with the cutaneous squamous cell carcinoma (cSCC) and non-malignant regions annotated. **d**, Spatial distribution of the mutation MT:12041:CC>TT, which is significantly enriched in the tumor region. **e**, Spatial distribution of the mutation MT:11872:C>T in non-malignant skin, which marks a spatial clone located near the skin surface and suggests a putative inward migration trajectory. **f**, For the cSCC1 sample, UMAP of transcriptional states from cSCC showing expression clusters. **g**, For the cSCC1 sample, spatial map of expression-based clusters in cSCC, with spots carrying MT:12041:CC>TT circled. **h**, For the cSCC1 sample, UMAP of transcriptional states from cSCC showing that MT:12041:CC>TT-positive spots are dispersed across expression space. **i**, For the Skin1 sample, UMAP of transcriptional states showing expression clusters in non-malignant skin. **j**, For the Skin1 sample, spatial map of expression-based clusters in non-malignant skin, with spots carrying MT:11872:C>T circled. **k**, For the Skin1 sample, UMAP of transcriptional states from non-malignant skin showing that MT:11872:C>T-positive spots are dispersed across expression space. **l**, For the cSCC1 sample, interferon pathways and migration-related pathways are enriched in the cSCC region. **m**, For the cSCC1 sample, comparison of MT:12041:CC>TT-positive spots with the remaining spots within the same non-tumor expression cluster, showing enrichment of interferon pathways in mutation-positive spots, consistent with possible cell migration.

Surprisingly, a similar spatial pattern was evident for the variant MT:11872:C>T described above in benign skin tissue (Skin1) (**Fig. 4e, Fig. S5c**). Pixels harboring this mutation were detected in the dermis, in addition to the clone’s dominant location in the skin epithelium. Moreover, some pixels in the dermis region containing this mutation were located outside hair follicles and secretory glands. This observation was unexpected because an intact basement membrane would ordinarily be expected to constrain such micro-migratory behavior of non-malignant cells, and inward migration of epidermal cells into the dermis in benign tissue has not been well documented. To further assess the generality of this phenomenon, we examined multiple skin samples and identified several additional mutations that exhibited the same spatial pattern, including MT:15298:C>T, 17:41582371:G>A, and X:12976989:C>T, in Skin1 and its biological replicate Skin2. Most of these mutations also showed UV-associated mutational patterns (**Fig. S6**). We also identified a similar event in another tumor sample, in which the mutation MT:5168:C>T was detected at the skin surface of histologically non-malignant tissue and showed a similar inward extension into the dermis (**Fig. S7a-c**). Importantly, these events could not be recovered from transcriptional profiles alone. Specifically, in these samples, pixels carrying MT:11872:C>T, MT:12041:CC>TT, and MT:5168:C>T were broadly dispersed across expression-based clusters and UMAP space rather than forming compact transcriptional groups (**Fig. 4f–k, Fig. S5d-e, Fig. S7d-e**).

To further support these recurrent migration events, we compared mutation-positive spots with the remaining spots within the same non-tumor expression cluster in cSCC1. Mutation-positive spots showed enrichment of interferon-α and interferon-γ response pathways, which were also enriched in tumor-associated spots relative to non-tumor spots (**Fig. 4l-m**). These findings were confirmed in the biological replicate sample cSCC2 (**Fig. S5b, S5d, S5f-g**). Given that both samples showed histological features of inflammation, local inflammation may further contribute to this phenomenon (**Fig. 4l–m, Fig. S5f–g**). Collectively, these observations indicate that mutation-informed spatial analysis by StarTrak can infer migratory events of mutant cells that would remain largely invisible to approaches based only on expression levels.

## Discussion

Understanding how tumor micro-evolution occurs within human tissues remains a central challenge in cancer biology. Although spatial genomics has significantly extended our ability to resolve tissue architecture with genomic data, the mutational information contained in these data remains largely unexploited. Here, we present StarTrak, a neural network framework that enables robust somatic variant detection coupled with faithful lineage reconstruction.

To overcome the sparsity of mutational information in spatial genomics datasets, StarTrak exploits iterative feedback between mutation detection and phylogenetic inference, jointly performing variant calling and lineage reconstruction within a unified framework. This approach reflects the intrinsic relationship between clones with distinct spatial patterns and the somatic mutations that define their identities: accurate lineage reconstruction refines *de novo* mutation calls within lineages, while reliable *de novo* mutation calls are essential for accurate lineage reconstruction. Moreover, StarTrak employs an all-evidence strategy that incorporates not only mitochondrial and nuclear genomic variants but also histopathological features, expression profiles, and spatial context into lineage inference to identify clones with distinct spatial patterns. The integration of histopathological features and expression profiles assumes that cells within the same clone are more likely, but not required, to share similar morphological features and expression profiles, respectively. This design is particularly critical for spatial genomics data, where pixel-level measurements often lack complete nucleotide coverage and may represent aggregates of multiple cells.

Our results demonstrate that StarTrak recovers biologically meaningful clonal architecture across both neoplastic and benign tissues, as evidenced by faithful reconstruction of experimentally defined lineages and well-established trajectories of hepatocarcinogenesis. Beyond lineage reconstruction, StarTrak enabled biological discoveries not apparent from expression-based analyses. In both non-malignant skin and cSCC, mutation-defined spatial patterns revealed migration-like behavior of mutant cells across tissue compartments. The identification of UV-associated variants, coupled with their absence in bulk DNA-seq of matched non-neoplastic tissue, strongly supported the somatic nature of these mutations and was consistent with UV-induced damage at the skin surface. Critically, mutation-bearing cells were spatially coherent yet transcriptionally dispersed, indicating that expression-level analyses alone cannot capture such migration events. This observation has broader implications: expression similarity is not always a reliable proxy for lineage relationships. More generally, these findings illustrate how spatially resolved mutation analysis can uncover dimensions of tissue dynamics that would remain obscured by transcriptomic profiling alone.

However, several limitations remain. First, most current spatial platforms do not achieve true single-cell resolution, and individual pixels often represent mixtures of cells with different phylogenetic origins. Consequently, spatial clonal reconstruction captures the phylogenetic history of the predominant cell population in each pixel rather than that of individual cells. Second, because mutational information in spatial genomics data is sparse, somatic mutation calling by StarTrak is not genome-wide and remains limited to genomic regions with sufficient sequencing depth. Third, phylogenetic inference in spatial settings may not uncover the full clonal diversity, as only a small fraction of true mutational events is observed. The resulting trees should therefore be interpreted as evidence-supported lineage models rather than definitive evolutionary histories.

Cross-tissue heterogeneity in mutational load represents a further limitation. Although we demonstrate the broad applicability of StarTrak across diverse sample types, including cSCC, pancreatic carcinoma, liver tumors, T-cell lymphoma, B-cell lymphoma, and benign skin, its performance remains strongly dependent on the underlying genomic variation within the input tissue. Specifically, mutational burden and the spectrum of genomic alterations can vary substantially across tissue types. For example, melanoma typically exhibits a high burden of single-nucleotide mutations, whereas glioblastoma is often characterized by more extensive copy number alterations. Therefore, the use of StarTrak should take into account the genomic landscape of the input tissue. Although StarTrak uses spatial information by default for somatic mutation calling and lineage reconstruction, it can also operate without spatial information. This mode may be particularly useful for cancers in which malignant cells exhibit substantial migration within the tissue, such as acute myeloid leukemia in the bone marrow, where extensive cell movement may diminish the utility of spatial distributions for mutation calling and lineage inference.

In summary, StarTrak establishes a general framework for spatially aware somatic variant detection and *in situ* lineage reconstruction across diverse spatial genomics platforms. By integrating mutational evidence with tissue context, StarTrak extends the utility of spatial genomics beyond describing molecular states to understanding evolutionary history within intact human tissues. It provides a foundation for investigating how clonal evolution, cell migration, and microenvironmental adaptation interact in both malignant and non-malignant tissues. We anticipate that this integrative strategy will prove broadly applicable to future spatial phylogenetic analyses.

## Methods

### Code availability

The StarTrak workflow was implemented in Bash and Python and requires a Linux operating system. Overall, the pipeline takes BAM files and associated metadata as input and performs comprehensive spatial analysis of RNA variants. Its source code is publicly available on GitHub at https://github.com/gersteinlab/StarTrak. A containerized version of the workflow is also provided in the same GitHub repository. Code for all figures in the manuscript is also available in this repository.

### Data availability

Spatial transcriptomic datasets used in this study included both publicly available data and datasets generated in-house. 10x Genomics Visium datasets and paired bulk DNA-sequencing data from human skin tissues were obtained from the Gene Expression Omnibus (GEO) under accession numbers GSE144240 and GSE173651 (*31, 32*). Additional 10x Genomics Visium datasets were obtained from the Human Tumor Atlas Network (*33*). Patho-DBiT and epi-Patho-DBiT datasets for MALT lymphoma were obtained from GEO under accession numbers GSE274641 and GSE288178, respectively (*13, 24*). Patho-DBiT datasets for AITL and HCC, as well as epi-Patho-DBiT datasets for AITL, will be made publicly available through separate studies.

### RNA variant matrix construction from BAM files

All sequencing datasets were mapped to the GRCh38 reference genome. Patho-DBiT datasets were processed by ASTRO, whereas other datasets were processed using 10x Genomics Space Ranger (*34, 35*). For each input BAM file, unmapped and supplementary alignments were removed. Multi-mapping and uniquely mapped reads were processed separately. For the standard curated BAM files, multi-mapping reads were removed. Reads containing skipped regions were split at those regions to prevent discontinuous alignments from contributing to variant calling.

Per-pixel variant information was then generated from the curated BAM files using samtools mpileup. Next, all variants were filtered based on allele frequency and read depth using counts summed across pixels. Variants were retained only if the pooled ALT count was at least 5, the total depth was at least 5, and the pooled VAF was at least 0.01. The resulting matrix was represented as a variant-by-pixel matrix, with variants as rows and pixels as columns. Variants were then annotated using the supplied single-nucleotide polymorphism (SNP) VCF file and RNA-editing VCF file to mark known SNPs and RNA-editing events. This annotated variant matrix was used for downstream analysis.

### Variant ranking for prioritizing potentially somatic mutations

VAF separation ranking applies filters across all variants **(Fig. S1a)**. Default requirements are a total read depth of at least 200, summed ALT and wild-type (WT) counts each exceeding 20, a pooled VAF of at least 0.01 (total ALT count divided by total read depth), and read depth of at least 2 in at least 1% of pixels. Subsequent separation scoring uses only pixels with a read depth of at least 3 at the variant site and requires at least 20 qualifying pixels per variant. The VAF separation ranking is calculated among covered pixels. For variant Vat pixel *i, a*_*vi*_ denotes the ALT read count, and *d*_*vi*_ denotes the total read depth. A pseudocount-adjusted VAF is calculated as follows:

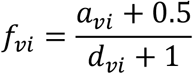

All covered pixels are then sorted by *f*_*vi*_ to identify an optimal cutoff of *f*_*vi*_, referred to as a split point, *k*, where k denotes the number of pixels assigned to the left side of the split (i.e., the split occurs between the k-th and (k+1)-th pixels in the *f*_*vi*_-ranked list). The mean VAFs on the left and right sides of the ordered list are computed as 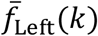 and 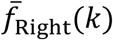 respectively, and used to calculate the split-specific VAF-rank separation score. Each split requires at least 10 pixels in both groups. The variant-level VAF-separation score, 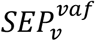, for variant Vis taken as the maximum score across all possible split points:

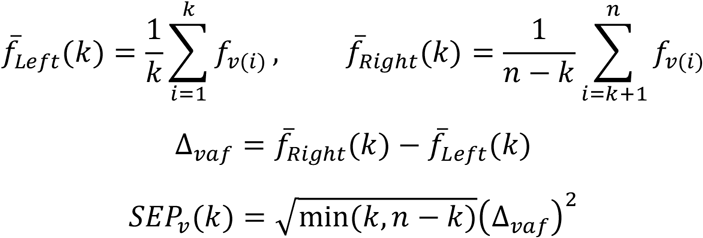

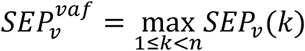

Here, *n* is the number of pixels covered at the variant, and *SEP*_*v*_*(k)* is a split-specific VAF-rank separation score.

Sufficient pixel coverage is required to evaluate the spatial distributions of reads. Therefore, spatial-distribution separation ranking is performed only for variants with ALT reads detected in at least 10 pixels and WT reads detected in at least 10 pixels. The remaining variants are retained for downstream analysis, with subsequent calculations restricted to the retained pixels for each corresponding variant. For each retained variant, spatial coordinates are normalized at the variant level as follows:

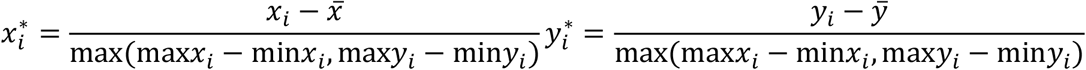

For variant Vat pixel *i, a*_*vi*_ denotes the ALT read count, *d*_*vi*_ denotes the total read depth, and *b*_*vi*_ = *d*_*vi*_ − *a*_*vi*_ denotes the observed WT count. The spatial distributions of ALT and WT reads across pixels are defined as:

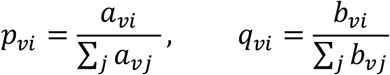

The spatial energy distance for variant Vis then defined as:

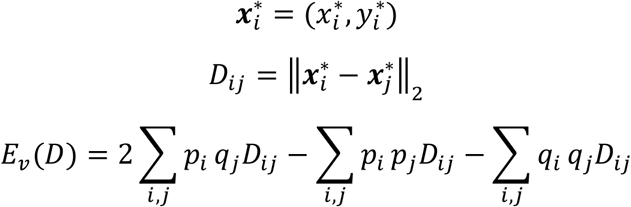

Equivalently,

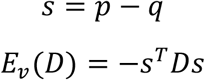

VAF separation ranking and spatial-distribution separation ranking are strongly influenced by per-pixel read depth. Therefore, a hypergeometric permutation procedure is used to estimate a variant-specific null distribution. For variant *v*, simulated ALT counts are sampled as follows:

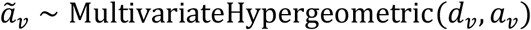

Here, *MultivariateHypergeometric* denotes the multivariate hypergeometric distribution; random samples from this distribution are generated in Python using ‘numpy.random.Generator.multivariate_hypergeometric’.

For each simulated replicate, the corresponding VAF separation score or spatial-distribution separation score is recalculated using the permuted ALT-count allocation. The mean and standard deviation are estimated across all simulated replicates, and the standardized score, *Z*_*v*_, is then defined as follows:

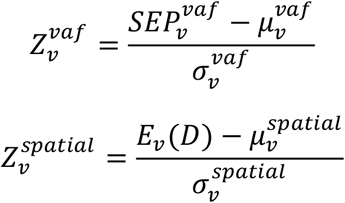

For spatial-distribution separation ranking, the raw energy distance also provides useful information. Therefore, the final spatial-distribution separation score, 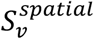, combines the calibrated z-score with the rank of the raw energy distance:

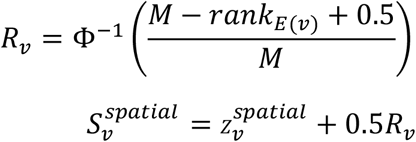

Here, *M* is the number of valid variants, *R*_*v*_is the rank-based inverse-normal score, and Φ^−1^ is the quantile function of the standard normal distribution.

### Multimodal embedding of pixels

StarTrak uses an autoencoder to learn pixel-level embeddings **(Fig. 1c)**. The encoder integrates available spatial coordinates, histopathological images, and gene expression profiles to generate embeddings for downstream phylogenetic reconstruction. For the expression modality, counts are log-normalized and up to 2,000 genes with the highest variance are selected. Their expression values are standardized to zero mean and unit variance and clipped to [−10, 10]. Principal component analysis is then applied to the resulting matrix. The input dimension of the expression modality is set to the smallest number of principal components explaining at least 60% of the total variance, capped at 40. Input variants are obtained from the two phylogeny-independent ranked variant lists described above, which are generated by ordering variants in descending order of 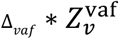and 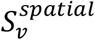, respectively. By default, the top 500 variants from each available ranking are combined and deduplicated to form the candidate pool. These candidates undergo quality-control filtering, and annotated SNPs and RNA-editing sites are excluded.

The autoencoder follows a standard multimodal architecture (**Fig. 1c)**. The training loss is the mean squared error across pixels and input features:

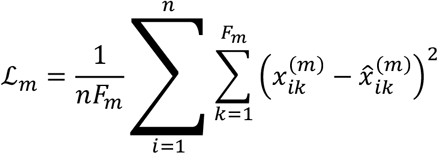

where *n* is the number of pixels, *F*_*m*_ is the number of input features in modality *m*, and 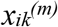 and 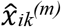 are the observed and reconstructed feature values.

### Iterative Tree Reconstruction with Candidate RNA Variant Updates

For phylogenetic reconstruction, StarTrak first defines a pool of putative somatic-like RNA variants using two tree-independent ranking strategies. Specifically, the top 500 variants from the spatial-distribution separation ranking and the top 500 variants from the VAF separation ranking are pooled and subjected to RNA variant quality control. From the quality control (QC)-passed candidate pool, StarTrak initializes tree construction with a compact seed set composed of the top 40 spatial-distribution-ranked variants and the top 20 variants from the VAF ranking. This seed set is used for initial phylogenetic tree construction. Details of the tree-building procedure, including the calculation of the pixel distance matrix, are described in the following section, “Embedding-Diffused Phylogenetic Tree Construction from RNA Variants.” The resulting tree is then used to evaluate additional candidate variants by their branch-level support. Variants passing the admission criteria are added to the admitted set, and the tree is rebuilt using the updated variant list. This iterative procedure continues until no additional candidates pass or the maximum number of iterations (set to 50 by default) is reached **(Fig. S1b)**.

### Embedding-Diffused Phylogenetic Tree Construction from RNA Variants

Considering the differences in ploidy between mitochondrial and nuclear genomic DNA, StarTrak applies smoothed VAFs for mitochondrial variants and binary VAFs for nuclear genomic variants during phylogenetic tree construction. For mitochondrial variants, the per-pixel VAF *VAF*_*vi*_ and evidence strength *e*_*vi*_ are:

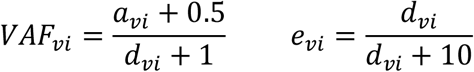

For non-mitochondrial variants, the per-pixel VAF and evidence strength are:

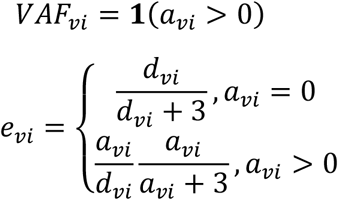

An evidence-weighted generalized Jaccard dissimilarity matrix (*D*_*ij*_) is then constructed for phylogenetic tree inference using average-linkage hierarchical clustering. The dissimilarity between two pixels is based on their per-variant VAF differences, with each variant contribution weighted by the evidence between the two pixels. Because mutational information is sparse at the individual-pixel level, StarTrak diffuses VAFs over the previous multimodal embedding before computing distances. Each embedding dimension is z-score standardized, and Euclidean distance is then used to identify the 12 most similar pixels for each pixel. These neighbors are weighted (*K*_*ij*_) by a Gaussian kernel, with closer pixels receiving larger weights. For each input variant, the initial VAFs and evidence are then diffused for two steps. At each step, each pixel gathers weighted averages from its embedding neighbors and updates its own state by mixing the previous value with the neighbor-averaged value:

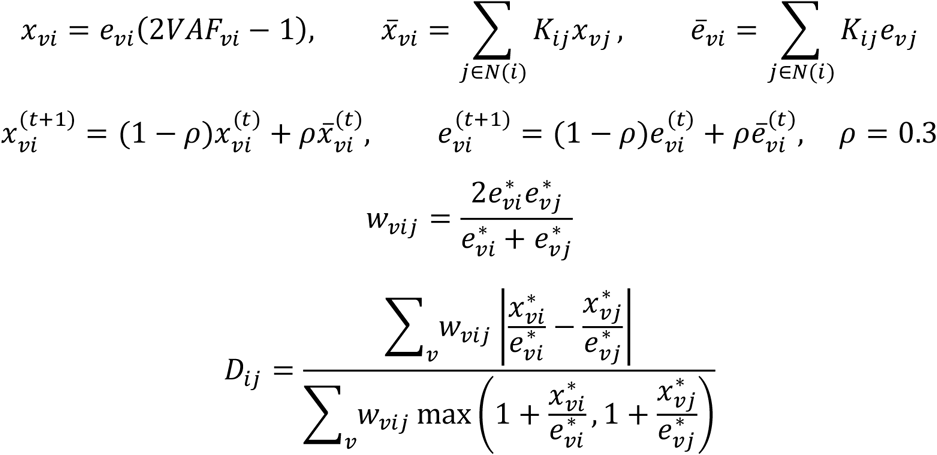

where ^*^ indicates the final value after diffusion.

### Tree-Guided Candidate RNA Variant Updating

After the initial tree is constructed, candidate RNA variants are re-evaluated against the current tree. For each candidate variant, branch-specific fit statistics are computed over all eligible tree branches. A candidate is retained only if it satisfies several constraints. The corresponding statistics are calculated as follows:

For each mitochondrial variant:

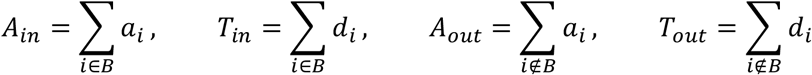

where B denotes the set of pixels contained in a specific subtree, A_in_ is the alternative-allele read count of the variant summed over B, T_in_ is the total read depth of the variant summed over B, A_out_ is the alternative-allele read count of the variant summed over all pixels not in B, and T_out_ is the total read depth of the variant summed over all pixels not in B.

For each non-mitochondrial variant:

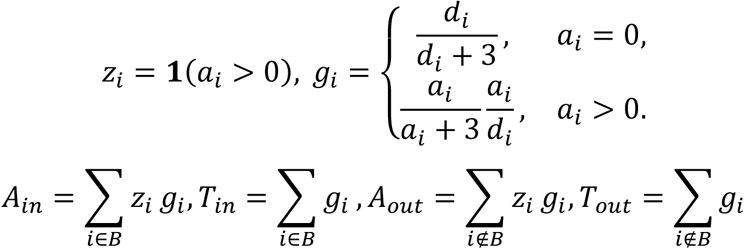

The branch-specific signal probabilities are:

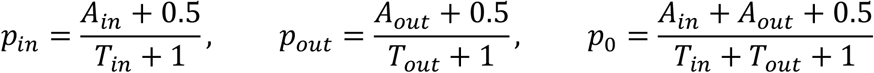

The branch-specific model is compared against a null model. For mitochondrial variants, StarTrak uses a binomial ALT/depth model:

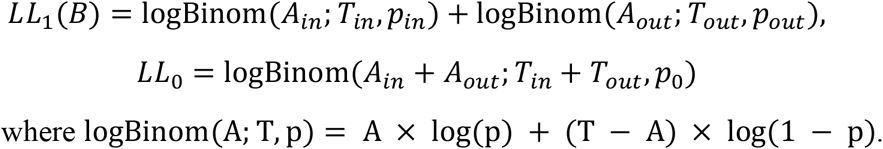

For non-mitochondrial variants, StarTrak uses a Poisson model:

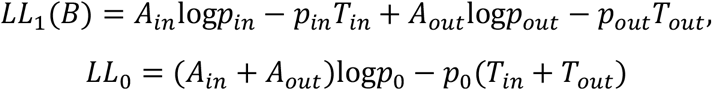

Among branches with T_in_ > 0, T_out_ > 0 and p_in_ > p_out_ (and A_in_ > 0 for non-mitochondrial variants), the best branch is selected by:

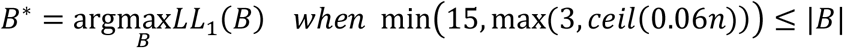

where |*B*| is the number of pixels in branch *B* and *n* is the total number of pixels.

Each candidate variant must also satisfy:

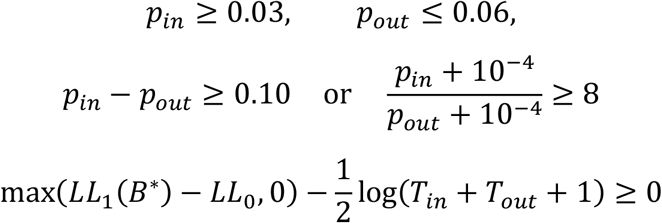

After candidate admission, the variant list is updated, the tree is rebuilt from the updated variant set, and the procedure is repeated until no additional candidate passes the admission criteria or the maximum number of update rounds is reached.

### Phylogeny-Guided Somatic Mutation Calling

After phylogenetic reconstruction, StarTrak performs somatic mutation calling. At this stage, only variants with ALT reads detected in at least 10 pixels and WT reads detected in at least 10 pixels are considered. Candidate variants are then evaluated through a series of biological and technical filters. This somatic mutation calling step additionally requires an initial tumor–normal pixel annotation, which can be performed manually based on paired histology images. To improve the robustness of this annotation, StarTrak refines pixel-level tumor–normal labels using the previously reconstructed phylogenetic tree. Each internal tree clade is examined as a candidate clonal compartment. For a clade (C) with at least 10 annotated pixels, the proportion of tumor-labeled pixels (*π*_*C*_) is calculated as follows:

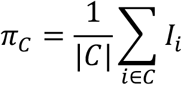

where *I*_*i*_ = 1 denotes a tumor-labeled pixel and *I*_*i*_ = 0 denotes a normal pixel.

If *π*_*C*_ ≥ 0.95, all pixels in the clade are assigned to the tumor compartment; otherwise, their original labels are retained. Refinement is performed once using the original labels, and originally tumor-labeled pixels remain in the tumor compartment. When a pixel belongs to multiple clades, membership in any qualifying clade is sufficient for tumor assignment.

For each candidate variant, ALT and WT counts are pooled separately across the refined tumor and normal compartments after limiting each pixel’s total depth contribution to 10. For pixels exceeding this depth, ALT and WT counts are scaled proportionally. Compartment-specific VAF is calculated as the pooled adjusted ALT count divided by the pooled adjusted total depth, and the tumor–normal VAF increase is defined as the tumor VAF minus the normal VAF. Candidates must have RNA coverage in both compartments and ALT support in at least three tumor pixels. Variants admitted to the phylogenetic tree require a tumor–normal VAF increase of at least 0.1. Variants not admitted to the tree require an increase of at least 0.25 and a 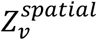 of at least 0.25. Raw calls are then subjected to read-level artifact filtering based on ALT base quality, mismatch bias, and ALT position bias, as described in the section “Quality control of RNA variants.” Variants passing these filters are written as the somatic call list.

### Quality control of RNA variants

StarTrak applies RNA variant quality control in two stages. First, annotation-based filtering is performed using supplied VCF files: variants annotated as known SNPs or RNA-editing sites are removed. For final somatic mutation calling, variants with a population allele frequency ≤0.01 remain eligible as somatic candidates. The reference datasets used in this manuscript for known SNPs and RNA-editing sites were obtained from gnomAD and REDIportal, respectively (*36, 37*). Second, RNA-specific quality filters are applied. Indels longer than two bases and reads with multiple mappings are removed. The filters also evaluate ALT base quality, mismatch bias, and ALT positional bias. Variants with a mean ALT base quality of 20 or lower are removed. Variants are also removed if ALT-supporting reads have a significantly higher adjusted mismatch burden than WT-supporting reads, defined by a one-sided Mann–Whitney U-test p-value of 0.0001 or lower and a median ALT–WT mismatch difference of at least 5. Variants are further removed if ALT-supporting reads are significantly closer to read ends, splice boundaries, or soft-clip boundaries (defined in the BAM CIGAR string) than WT-supporting reads, defined by a one-sided Mann–Whitney U-test p-value of 0.0001 or lower, a median ALT distance to the nearest boundary of 5 bp or less, and a median WT–ALT distance difference of at least 5 bp. For FFPE samples, a strand-aware FFPE artifact filter is applied to C>T candidate variants. A variant is removed as a putative FFPE artifact if at least 90% of its ALT-supporting reads are represented as C>T changes on the RNA strand and if either its VAF is below 0.25 or its ALT depth is below 3. For data from the DBiT platform, variants are excluded if the ratio of the larger to the smaller number of distinct ALT-supporting coordinates along the two spatial axes exceeds 10. This filter targets variants with highly elongated, strip-like spatial support patterns that may reflect technical artifacts.

### Benchmarking of somatic mutation calling

Somatic mutation call benchmarking compares variant calls from StarTrak, Monopogen and SComatic using matched bulk DNA sequencing data as the ground truth. For each candidate variant, allele counts are extracted from the corresponding tumor and normal bulk DNA BAM files. A candidate is labeled “somatic” if the tumor allele fraction is greater than the normal allele fraction. If the tumor allele fraction is not greater than the normal allele fraction, the candidate is labeled “non-somatic” only when the tumor WT read count is at least 10 and the normal WT read count is at least 10. All other RNA variants are excluded. This rule defines positive labels by tumor enrichment rather than by strict somatic calls from a specialized bulk DNA somatic variant caller, because the sequencing depth of the matched bulk DNA samples is limited. On the RNA side, a variant is included in the benchmark only if it is detected in at least 10 ALT-positive pixels and at least 10 WT-positive pixels.

Precision for each method is defined as the proportion of called variants that are labeled “somatic” based on matched bulk DNA evidence. To calculate recall, we first take the union of variants called by the three methods. The recall denominator is defined as the total number of variants in this union that are labeled “somatic.” For each method, the recall numerator is the number of its called variants that are labeled “somatic.” Recall is therefore calculated as the fraction of pooled putative somatic variants recovered by each method. All three methods were evaluated within a common candidate set, excluding multimapping variants, RNA-editing sites, high-frequency SNPs and variants with recorded QC failures.

### Benchmarking lineage reconstruction with experiment-based ground truth

To evaluate StarTrak’s performance in lineage reconstruction, we utilized three single-cell datasets generated with experimentally defined clonal identities as ground truth. The benchmark datasets included one dataset from the A549 cell line and two datasets from the TF-1 cell line (*12, 27*). Because some small clones contained too few cells for robust evaluation, clone subsets were selected for benchmarking: the two largest clones were included for the A549 dataset, and the four largest clones were used in the second TF-1 dataset. All clones were used for the first TF-1 dataset. For each dataset, single cells were processed using the single-cell mode of the StarTrak pipeline (**Supplementary Material**). The lineage structure inferred by StarTrak was then compared with the ground-truth lineage labels. Agreement between clone labels and the inferred phylogenetic tree was quantified using accuracy, which was computed by assigning each ground-truth clone to at most one monophyletic clade in the tree. Specifically, the algorithm searched for the valid set of non-overlapping clades that maximized the total number of correctly assigned labeled cells.

Accuracy was then calculated as this maximum number of correctly assigned cells divided by the total number of cells with ground-truth labels. For visualization, trees were plotted using ggtree (v3.10.0) (*38*).

## Author contributions

Project conceptualization: D.Z., M.G., Jun L., and R.F.; data curation: D.Z., Y.J., Y.H., J.T., Jiaqi L., and D.B.; data analysis: D.Z., Y.J., X.Y., Y.H., D.B., and Z.C.; funding acquisition: M.G., R.F. and Jun L.; methodology design and development: D.Z., X.Y., Y.J., D.B., M.G., and Jun L.; experimental data provided by: R.F., Z.B., A.E., X.T., and H.L.; scientific discussion: D.Z., N.Y., Y.C., J.T., and Jiaqi L.; writing: D.Z., Jun L., and M.G.

