## Supplementary figures for "Determining cellular phylogeny and migration at high resolution in spatial genomics"

### Supplementary Methods:

##### Application of StarTrak to single-cell datasets

For single-cell datasets without spatial measurements, candidate variants are ranked using variant allele fraction (VAF) separation scores. Normalized expression profiles are first reduced by principal component analysis (PCA) and then combined with mutation-profile inputs in the autoencoder. The resulting embedding defines the cellular neighborhood graph used for lineage reconstruction.

##### Spatial analysis of RNA variants from multiple-mapping reads

StarTrak also provides an optional module for analyzing variants supported by multiple-mapping reads. In this procedure, an additional binary alignment map (BAM) file containing all multiple-mapping reads is generated. During construction of the corresponding RNA variant matrix from this BAM file, one representative alignment is retained for each multiple-mapping read group. By default, this additional RNA variant matrix is used only to identify spatially informative candidate variants and is not included in downstream lineage reconstruction. Genomic-source inference is then performed for each such multiple-mapping RNA variant.

Candidate gene sources are inferred for variants supported by multimapping reads. For each variant, alternative allele (ALT)-supporting read names are identified, and all associated alignments in the input BAM file are examined for overlap with exons annotated in the reference gene transfer format (GTF) file. Candidate genes are pooled across alignments for each read name. Each read name associated with k candidate genes contributes a weight of 1/k to each gene. These weights are summed across ALT-supporting read names to obtain a gene-assignment support score for each variant–gene pair.

##### Comparison of mitochondrial and nuclear sequencing depth

Mean sequencing depth was calculated separately for the nuclear and mitochondrial genomes in each BAM file using samtools depth. Chromosome Y was excluded to avoid sex-related differences in sequencing coverage. For each sample, mean nuclear depth was obtained by summing per-base read depths across the included nuclear chromosomes and normalizing the total to their combined reference length. Mean mitochondrial depth was calculated analogously using the mitochondrial genome length. Examining 25 spatial transcriptomic datasets (including both Patho-DBiT (*1*) and 10x Visium datasets) and 11 spatial ATAC-seq (epi-Patho-DBiT) datasets (*2*), we found that mitochondrial reads were abundant in both data types, reaching mean read coverage of 30.2 to 100,560.5 in spatial transcriptomic data and 8,144.1 to 174,164.3 in spatial ATAC-seq data. In contrast, mean read coverage in the nuclear genomes (0.02 to 4.88) was much lower for both data types.

**Identification of high-confidence somatic variants using bulk DNA sequencing data**

Bulk DNA–derived high-confidence somatic variant lists were generated using cohort-specific strategies according to the availability of matched normal samples. For datasets with matched tumor and normal DNA, somatic single-nucleotide variants (SNVs) were identified using Strelka (v2.9.10). PASS variants were retained if they were supported by at least five ALT reads in the tumor, had a tumor VAF of at least 3%, had a normal VAF of no more than 1.5%, and had a normal VAF less than 30% of the corresponding tumor VAF. Non-PASS variants with strong tumor-specific support were also retained if they were supported by at least ten ALT reads and had a VAF of at least 30% in the tumor, had no ALT-supporting reads in the matched normal sample, and had a normal sequencing depth of at least 30. For datasets without matched normal DNA, variants were called from tumor DNA using GATK Mutect2 in tumor-only mode with a panel of normals and the gnomAD germline resource. Calls were processed using FilterMutectCalls, and only PASS variants with a sequencing depth of at least 20, at least five ALT-supporting reads, and a VAF of at least 5% were retained.

**Comparison of spatial ATAC-seq and spatial RNA-seq**

Sequencing coverage and variant-detection capacity were compared between spatial ATAC-seq datasets from epi-Patho-DBiT and spatial RNA sequencing (RNA-seq) datasets from 10x Visium and Patho-DBiT. Per-base sequencing depth was calculated from each BAM file using samtools depth. For each chromosome and dataset, the coverage fraction at a specified depth threshold was defined as the proportion of reference bases with a read depth greater than or equal to that threshold. Coverage fractions were calculated separately at depth thresholds of 2 and 5 and displayed on a logarithmic scale.

For high-confidence somatic variants, a variant was considered covered in a spot if at least one ALT read was detected. For each matched patient, a shared variant set was defined as the union of variants covered in at least one spatial dataset from that patient. All variants in this shared set were considered for visualization in the density plots. Two complementary metrics were calculated: the number of variants covered per spot and the number of spots covering each variant.

Concordance with matched DNA sequencing data was assessed for variants detected in the spatial datasets. Candidate loci were examined in the corresponding DNA-seq BAM files using samtools mpileup. Variants were considered eligible if they were supported by more than 10 ALT reads in the spatial dataset and had a total DNA-seq depth greater than 20. An eligible variant was considered confirmed if at least one DNA-seq read supported the ALT allele. For each spatial dataset, the confirmation rate was calculated as the proportion of eligible variants confirmed by the matched DNA sequencing data. Confirmation rates were compared between spatial ATAC-seq and spatial RNA-seq datasets using a two-sided Mann–Whitney U test.

**Mutation-associated gene expression analysis**

Gene expression profiles from four samples (Skin1, Skin2, cSCC1, and cSCC2) were analyzed using Seurat. Expression counts were normalized, centered, and scaled before PCA. The number of principal components retained for downstream analysis was determined from elbow plots, after which clustering was performed using FindClusters. The mitochondrial variant MT:11872:C>T was examined in Skin1 and Skin2, whereas MT:12041:CC>TT was examined in cSCC1 and cSCC2. Mutation-positive spots were defined as those containing at least one ALT read and having a VAF of at least 0.50. These spots were visualized in uniform manifold approximation and projection space and on spatial cluster maps.

Differential expression analysis was performed in cSCC1 and cSCC2 using FindMarkers. The first comparison was between manually annotated tumor-associated spots and non-tumor spots. The second comparison was performed within each cluster containing a sufficient number of mutation-positive spots, contrasting spots with a VAF more than 0.02 against the remaining spots in the same cluster. Genes were ranked by average log2 fold change and subjected to gene set enrichment analysis using fgsea (v1.28.0). The Hallmark and Gene Ontology Biological Process gene sets used in this analysis were obtained using msigdbr (v26.1.0).

**Mutation-order inference**

Selected variants are mapped onto the reconstructed lineage tree using spot-level ALT and total read counts. Variant placement is evaluated using the same statistical models as in the tree-admission stage. Uncertainty in node placement is assessed by resampling spots with replacement while holding the reconstructed tree fixed. Variant placement is repeated for each of 200 bootstrap replicates, and node support is defined as the proportion of replicates assigning a variant to a given node. The putative relative order of mutation acquisition is then inferred from the phylogenetic relationships among the assigned nodes.

##### Inference of cell migration from spatial variant distributions

SNVs within the top 0.5% of both $E_v(D)$ and $Z_v^{spatial}$ are selected. Dominant mutational regions are delineated by Gaussian smoothing of spot-level VAF, followed by thresholding using Otsu's method. For each variant, up to the two largest spatially connected components of threshold-positive spots are retained as mutational regions. Variants are included in subsequent testing only if these regions together contain at least 80% of threshold-positive spots. Outside the mutational regions, a one-sided exact binomial test assesses whether target substitutions occur above background levels by comparing the target ALT count with the combined counts of the other two non-reference bases (only base observations with Phred quality ≥25 are included). Variants with adjusted P values < 0.05 are considered markers of potential migration events.

##### Variant nomenclature

This table maps the variant labels used in the manuscript and figures to their corresponding Human Genome Variation Society (HGVS) descriptions:

| Variant label used in the manuscript and figures | HGVS description |
|---|---|
| MT:1791:G>A | NC_012920.1:m.1791G>A |
| MT:2069:T>C | NC_012920.1:m.2069T>C |
| MT:3608:G>A | NC_012920.1:m.3608G>A |
| MT:5168:C>T | NC_012920.1:m.5168C>T |
| MT:4569:G>A | NC_012920.1:m.4569G>A |
| MT:7527:G>A | NC_012920.1:m.7527G>A |

| | |
|---|---|
| MT:7608:G>A | NC_012920.1:m.7608G>A |
| MT:8276:C>T | NC_012920.1:m.8276C>T |
| MT:11872:C>T | NC_012920.1:m.11872C>T |
| MT:12041:CC>TT | NC_012920.1:m.12041_12042delinsTT |
| MT:15298:C>T | NC_012920.1:m.15298C>T |
| 2:210069871:C>G | NC_000002.12:g.210069871C>G |
| 4:1707216:A>C | NC_000004.12:g.1707216A>C |
| 7:107770272:T>C | NC_000007.14:g.107770272T>C |
| 11:65499155:T>A | NC_000011.10:g.65499155T>A |
| 11:65499746:G>A | NC_000011.10:g.65499746G>A |
| 11:65503603:T>A | NC_000011.10:g.65503603T>A |
| 14:24249195:C>T | NC_000014.9:g.24249195C>T |
| 17:41582371:G>A | NC_000017.11:g.41582371G>A |
| 19:18575179:C>T | NC_000019.10:g.18575179C>T |
| X:12976989:C>T | NC_000023.11:g.12976989C>T |
| X:154531643:C>T | NC_000023.11:g.154531643C>T |

(Note: the two adjacent SNVs comprising MT:12041:CC>TT were highly correlated (correlation coefficient = 0.99) and were treated as a single mutation marker in downstream analyses.)

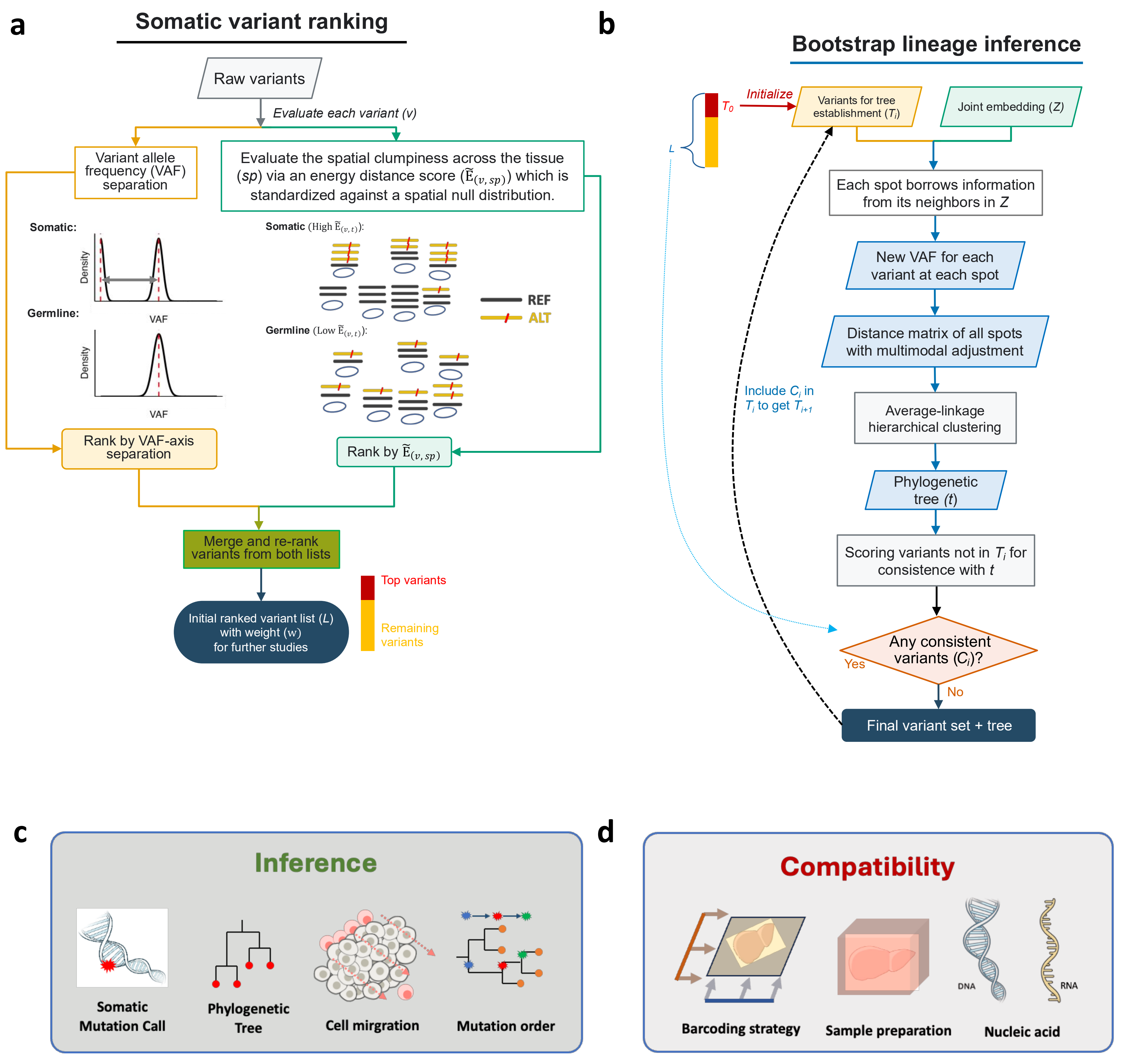
a
Somatic variant ranking
Raw variants
Evaluate each variant (v)
Variant allele frequency (VAF) separation
Evaluate the spatial clumpiness across the tissue (sp) via an energy distance score ($\tilde{E}_{(v,sp)}$) which is standardized against a spatial null distribution.
Somatic:
Density
VAF
Germline:
Density
VAF
Somatic (High $\tilde{E}_{(v,t)}$):
Germline (Low $\tilde{E}_{(v,t)}$):
REF
ALT
Rank by VAF-axis separation
Rank by $\tilde{E}_{(v,sp)}$
Merge and re-rank variants from both lists
Initial ranked variant list (L) with weight (w) for further studies
Top variants
Remaining variants
b
Bootstrap lineage inference
L
$T_0$
Initialize
Variants for tree establishment ($T_i$)
Joint embedding (Z)
Each spot borrows information from its neighbors in Z
New VAF for each variant at each spot
Distance matrix of all spots with multimodal adjustment
Include $C_i$ in $T_i$ to get $T_{i+1}$
Average-linkage hierarchical clustering
Phylogenetic tree (t)
Scoring variants not in $T_i$ for consistence with t
Any consistent variants ($C_i$)?
Yes
No
Final variant set + tree
c
Inference
Somatic Mutation Call
Phylogenetic Tree
Cell mirgration
Mutation order
d
Compatibility
Barcoding strategy
Sample preparation
DNA
RNA
Nucleic acid

Figure S1

**Figure S1. Architecture of the major stages of StarTrak. a,** Somatic variant ranking workflow. Candidate variants are ranked by VAF separation and spatial read-stack clumpiness to generate a ranked variant list. **b,** Phylogenetic tree construction. Candidate RNA variants are transformed into spot-level variant features and used to construct an average-linkage phylogenetic tree. The inferred tree is then used to update the candidate RNA variant set, and the updated variant set is used to rebuild the tree iteratively until convergence. **c,** StarTrak supports multiple analyses, including somatic variant calling, phylogenetic reconstruction, cell migration inference and mutation order inference. **d,** StarTrak is broadly compatible across sample preservation methods (FFPE and fresh-frozen specimens), spatial transcriptomics platforms (DBiT-seq and 10x Genomics Visium) and molecular types (RNA and DNA).

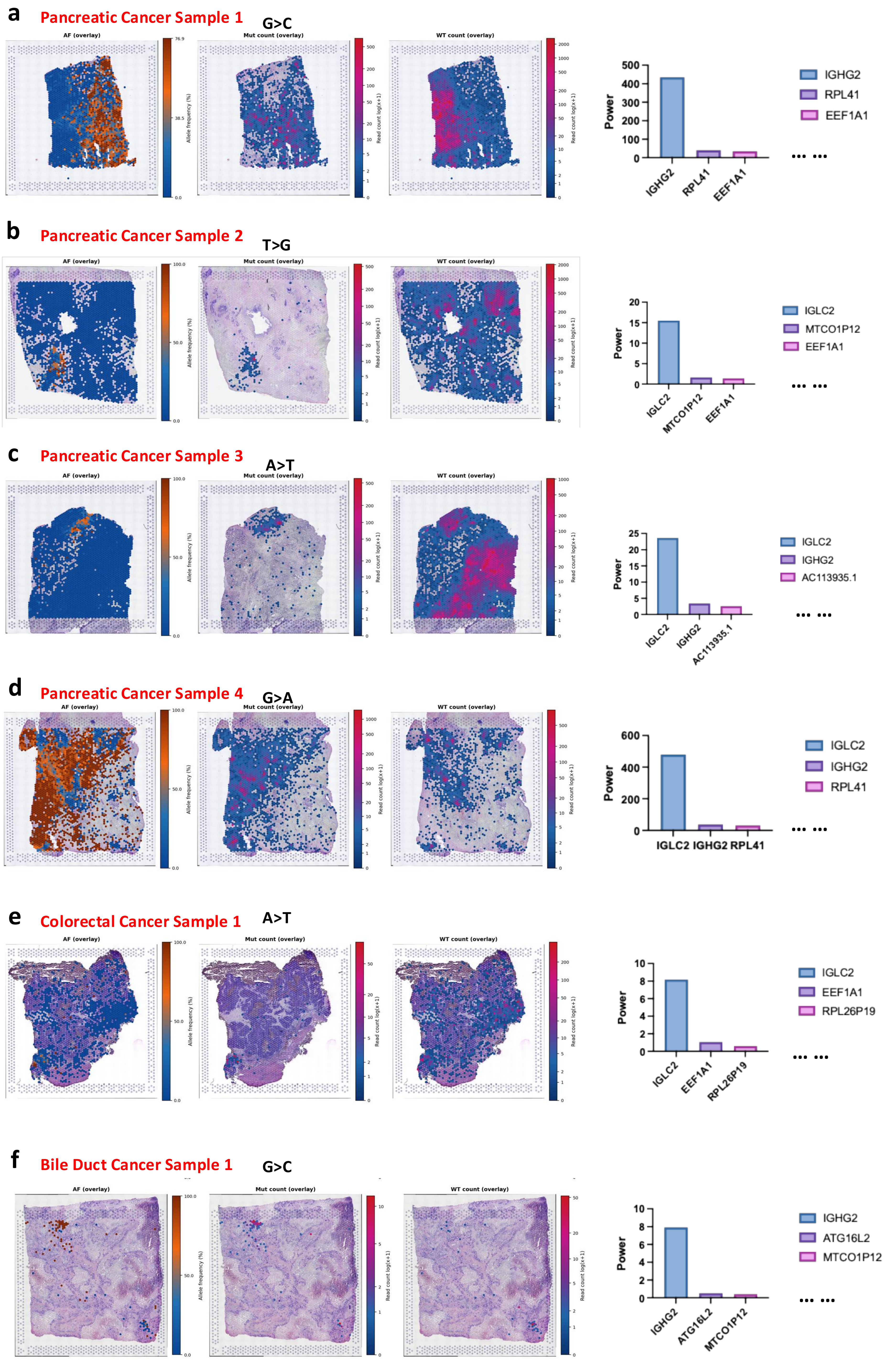
a
Pancreatic Cancer Sample 1
G>C
AF (overlay)
Mut count (overlay)
WT count (overlay)
Allele frequency (%)
Read count log(x+1)
Power
IGHG2
RPL41
EEF1A1
... ...
b
Pancreatic Cancer Sample 2
T>G
IGLC2
MTCO1P12
EEF1A1
c
Pancreatic Cancer Sample 3
A>T
IGLC2
IGHG2
AC113935.1
d
Pancreatic Cancer Sample 4
G>A
IGLC2
IGHG2
RPL41
e
Colorectal Cancer Sample 1
A>T
IGLC2
EEF1A1
RPL26P19
f
Bile Duct Cancer Sample 1
G>C
IGHG2
ATG16L2
MTCO1P12

Figure S2

**Figure S2. Characterization of variants with spatial patterns supported by multi-mapping reads across cancer samples.** Representative variants are shown for pancreatic cancer **a-d**, colorectal cancer **e**, and bile duct cancer **f**. For each example, the spatial distributions of allele fraction, mutant-read count, and reference-read count are overlaid on the tissue image, while the bar plot shows gene-assignment weights for each gene.

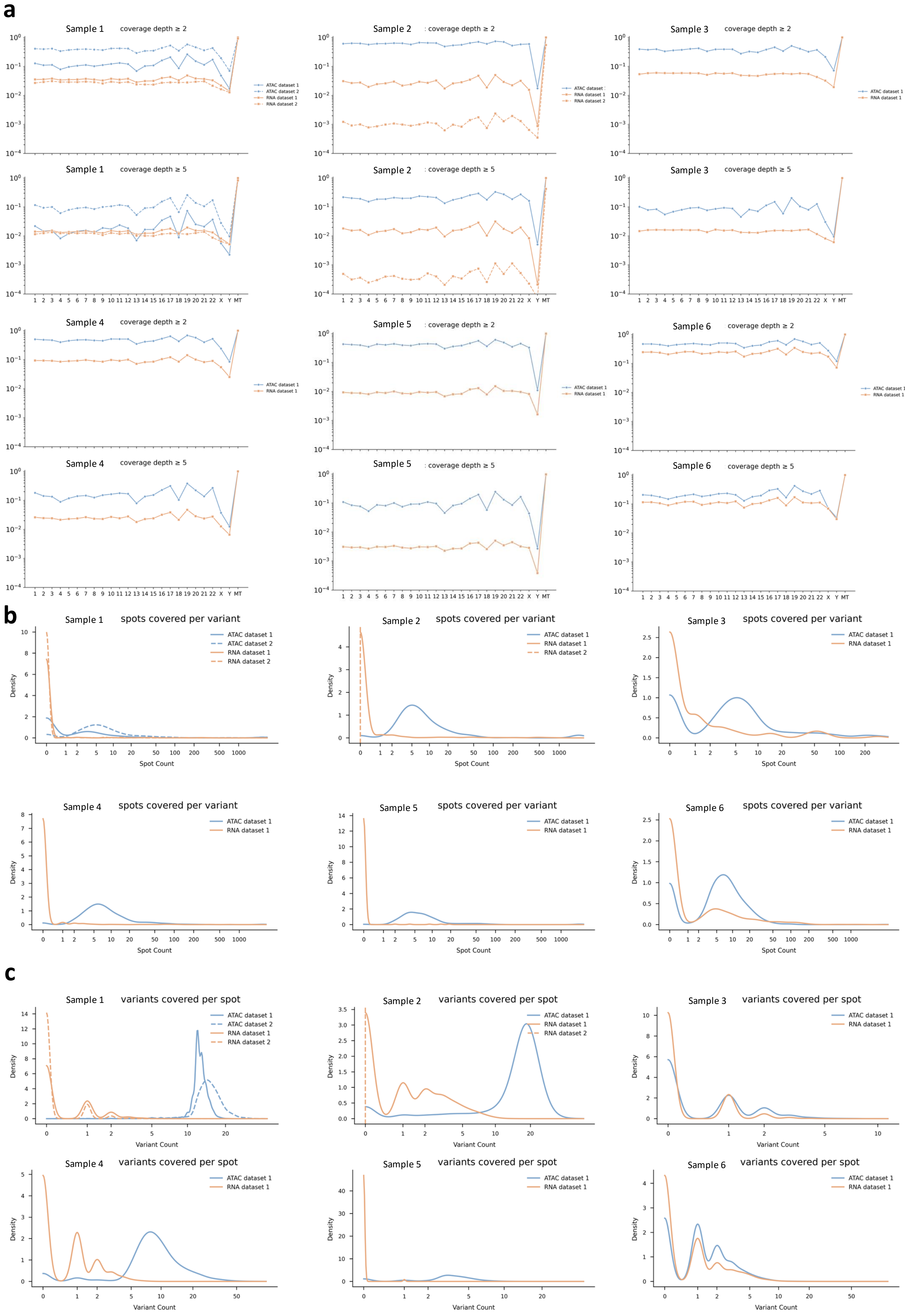

a
Sample 1 coverage depth ≥ 2
Sample 2 : coverage depth ≥ 2
Sample 3 coverage depth ≥ 2
ATAC dataset 1
ATAC dataset 2
RNA dataset 1
RNA dataset 2
Sample 1 coverage depth ≥ 5
Sample 2 : coverage depth ≥ 5
Sample 3 coverage depth ≥ 5
Sample 4 coverage depth ≥ 2
Sample 5 : coverage depth ≥ 2
Sample 6 coverage depth ≥ 2
Sample 4 coverage depth ≥ 5
Sample 5 : coverage depth ≥ 5
Sample 6 coverage depth ≥ 5
1 2 3 4 5 6 7 8 9 10 11 12 13 14 15 16 17 18 19 20 21 22 X Y MT
b
Sample 1 spots covered per variant
Sample 2 spots covered per variant
Sample 3 spots covered per variant
Sample 4 spots covered per variant
Sample 5 spots covered per variant
Sample 6 spots covered per variant
Density
Spot Count
c
Sample 1 variants covered per spot
Sample 2 variants covered per spot
Sample 3 variants covered per spot
Sample 4 variants covered per spot
Sample 5 variants covered per spot
Sample 6 variants covered per spot
Density
Variant Count


**Figure S3**

**Figure S3. Comparison of spatial ATAC-seq and spatial RNA-seq for genomic coverage and variant detection.** Spatial ATAC-seq and spatial RNA-seq were represented by epi-Patho-DBiT and Patho-DBiT, respectively. Each panel compares paired datasets from the same sample across different performance metrics: **a,** Genome-wide coverage. **b,** Number of spots covered per variant. **c,** Number of variants detected per spot.

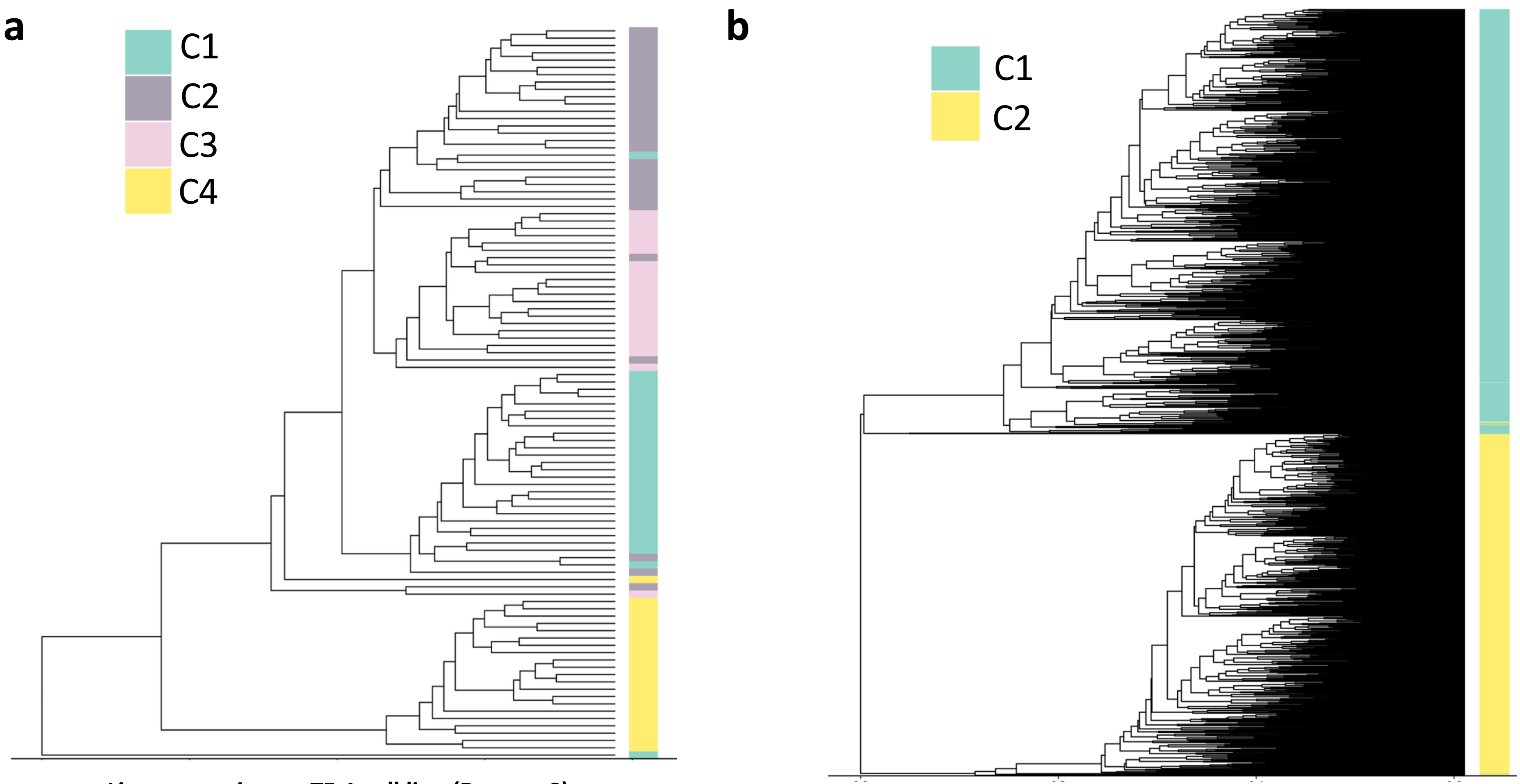


**Lineage tracing on TF-1 cell line (Dataset 2)**

**Lineage tracing on A549 cell line**

**Figure S4**

**Figure S4. StarTrak works for lineage reconstruction. a,** StarTrak recovers ground-truth clonal structures in another TF-1 cell-line dataset with experimentally defined ground truth. **b,** StarTrak recovers ground-truth clonal structures in an A549 cell-line dataset with CRISPR–Cas9-based lineage tracing.

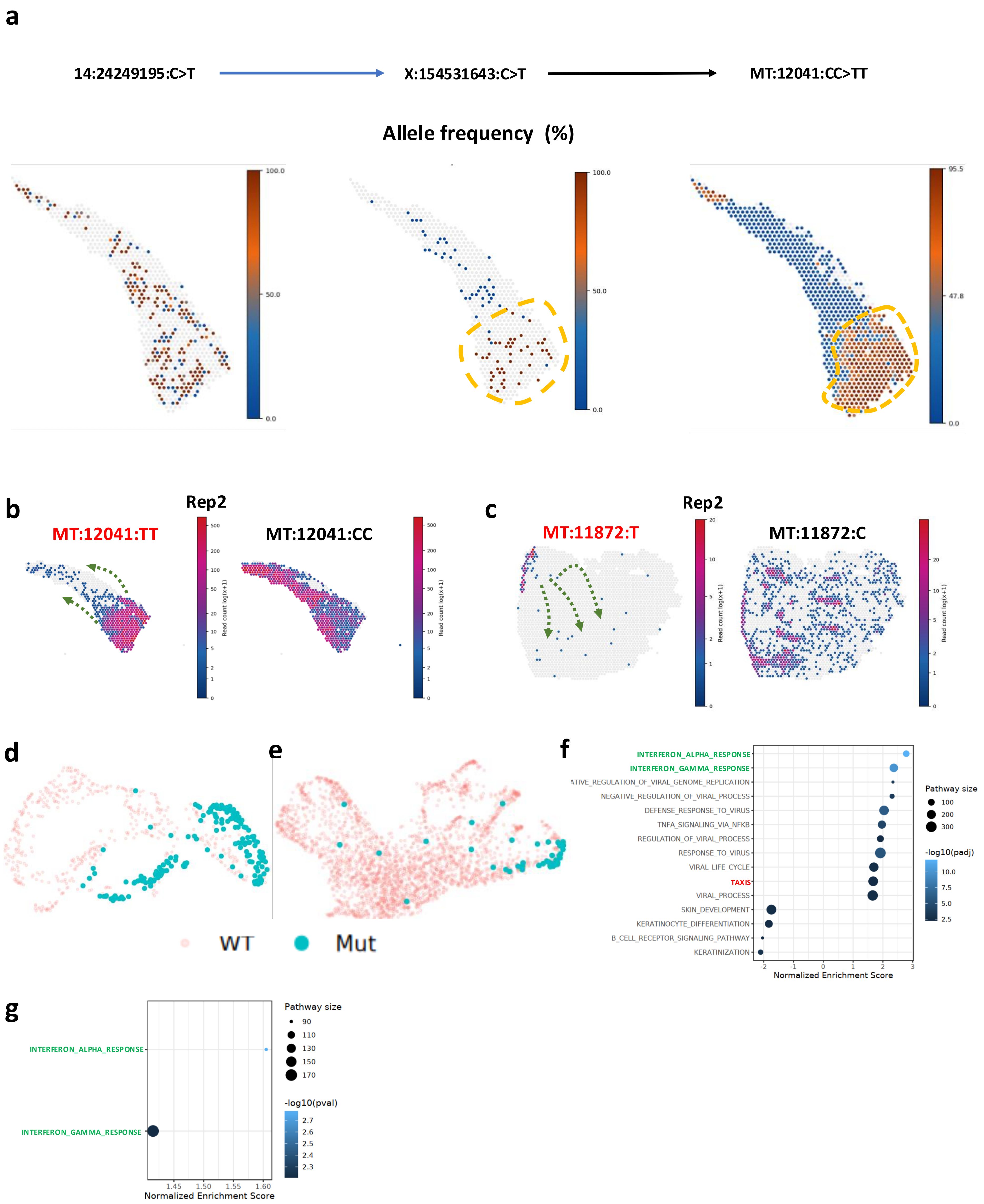

a
14:24249195:C>T
X:154531643:C>T
MT:12041:CC>TT
Allele frequency (%)
100.0
50.0
0.0
95.5
47.8
b
Rep2
MT:12041:TT
MT:12041:CC
Read count log(x+1)
c
Rep2
MT:11872:T
MT:11872:C
d
e
WT
Mut
f
INTERFERON_ALPHA_RESPONSE
INTERFERON_GAMMA_RESPONSE
NEGATIVE_REGULATION_OF_VIRAL_PROCESS
DEFENSE_RESPONSE_TO_VIRUS
TNFA_SIGNALING_VIA_NFKB
REGULATION_OF_VIRAL_PROCESS
RESPONSE_TO_VIRUS
VIRAL_LIFE_CYCLE
TAXIS
VIRAL_PROCESS
SKIN_DEVELOPMENT
KERATINOCYTE_DIFFERENTIATION
B_CELL_RECEPTOR_SIGNALING_PATHWAY
KERATINIZATION
Normalized Enrichment Score
Pathway size
-log10(padj)
g
INTERFERON_ALPHA_RESPONSE
INTERFERON_GAMMA_RESPONSE
Normalized Enrichment Score
Pathway size
-log10(pval)


**Figure S5**

**Figure S5. Spatial analysis of a biological replicate supports previously unrecognized migration of mutation-bearing cells in skin tissue. a,** StarTrak inferred the sequential acquisition of three mutational events: 14:24249195:C>T, X:154531643:C>T, and MT:12041:CC>TT. **b,** Spatial distribution of MT:12041:CC>TT in a biological replicate of the cSCC sample, showing significant enrichment of this mutation in the tumor region. **c,** Spatial distribution of MT:11872:C>T in a biological replicate of the normal skin sample. This mutation marks a spatial clone located near the skin surface and suggests a putative inward migration trajectory. **d,** For the cSCC2 sample, UMAP of transcriptional states from cSCC, showing that MT:12041:CC>TT-positive spots are dispersed across expression space in the biological replicate shown in **b**. **e,** For the Skin2 sample, UMAP of transcriptional states from normal skin, showing that MT:11872:C>T-positive spots are dispersed across expression space in the biological replicate shown in **c**. **f,** For the cSCC2 sample, interferon and migration-related pathways are enriched in the cSCC region in the biological replicate shown in **b**. **g,** For the cSCC2 sample, comparison of MT:12041:CC>TT-positive spots with the remaining spots within the same non-tumor expression cluster, showing enrichment of interferon pathways in mutation-positive spots.

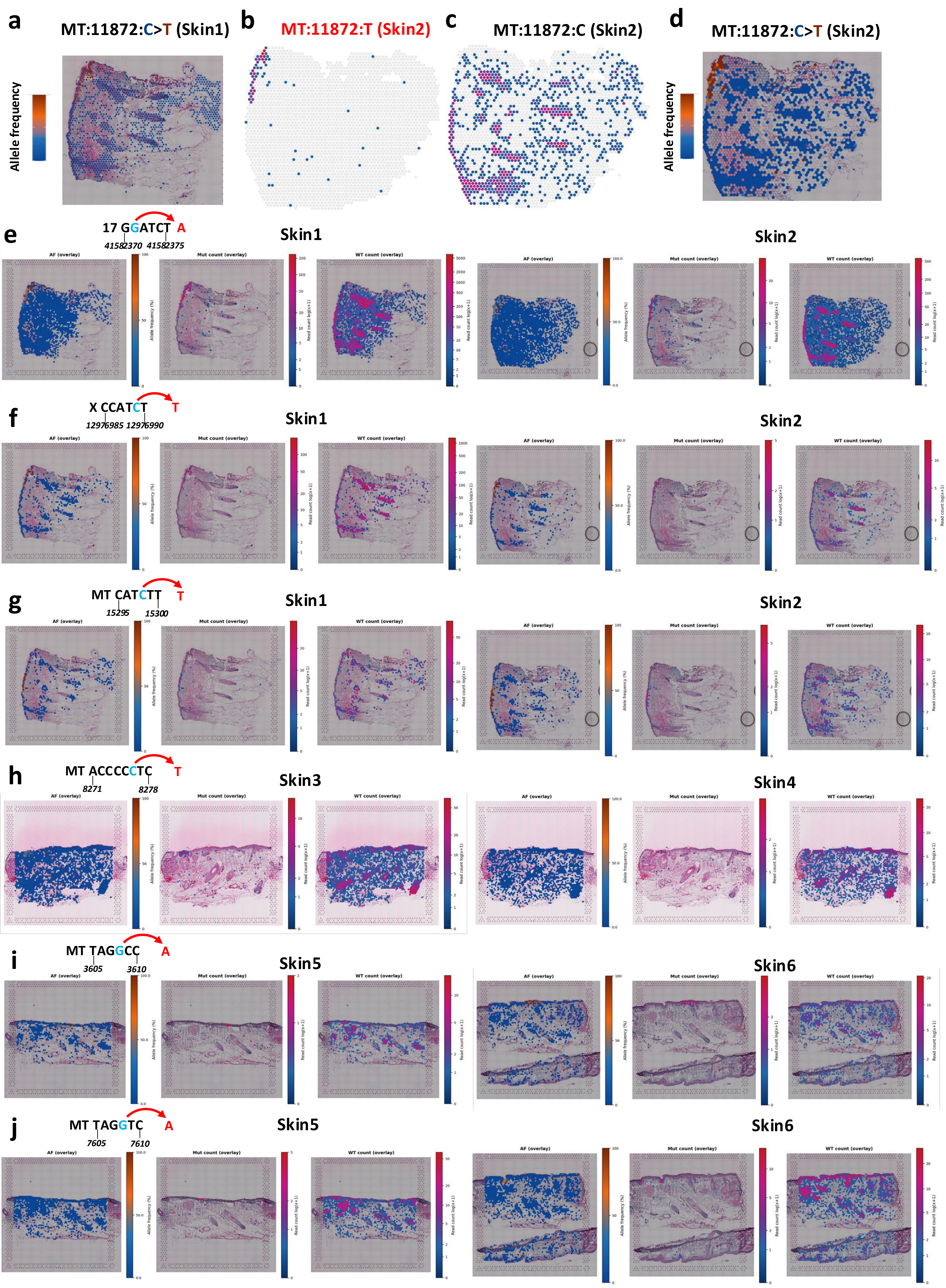

a
MT:11872:C>T (Skin1)
Allele frequency
b
MT:11872:T (Skin2)
c
MT:11872:C (Skin2)
d
MT:11872:C>T (Skin2)
Allele frequency
e
17 GGATCT A
41582370 41582375
Skin1
Skin2
f
X CCATCT T
12976985 12976990
Skin1
Skin2
g
MT CATCTT T
15295 15300
Skin1
Skin2
h
MT ACCCCCTC T
8271 8278
Skin3
Skin4
i
MT TAGGCC A
3605 3610
Skin5
Skin6
j
MT TAGGTC A
7605 7610
Skin5
Skin6


**Figure S6**

**Figure S6**. **Spatial patterns of UV-associated mutations across multiple skin samples.** **a,** Spatial distribution of the variant allele frequency of MT:11872:C>T in normal skin. **b-d,** Spatial distributions of the mutant allele, the reference allele, and the variant allele frequency of MT:11872:C>T in a biological replicate of the normal skin sample shown in **a**. **e,** Spatial distribution of 17:41582371:G>A in normal skin samples shown in **a-b**. **f,** Spatial distribution of X:12976989:C>T in normal skin samples shown in **a-b**. **g,** Spatial distribution of MT:15298:C>T in normal skin samples shown in **a-b**. **h,** Spatial distribution of MT:8276:C>T in normal skin samples with a biological replicate. **i,** Spatial distribution of MT:3608:G>A in normal skin samples with a biological replicate. **j,** Spatial distribution of MT:7608:G>A in normal skin samples with a biological replicate.

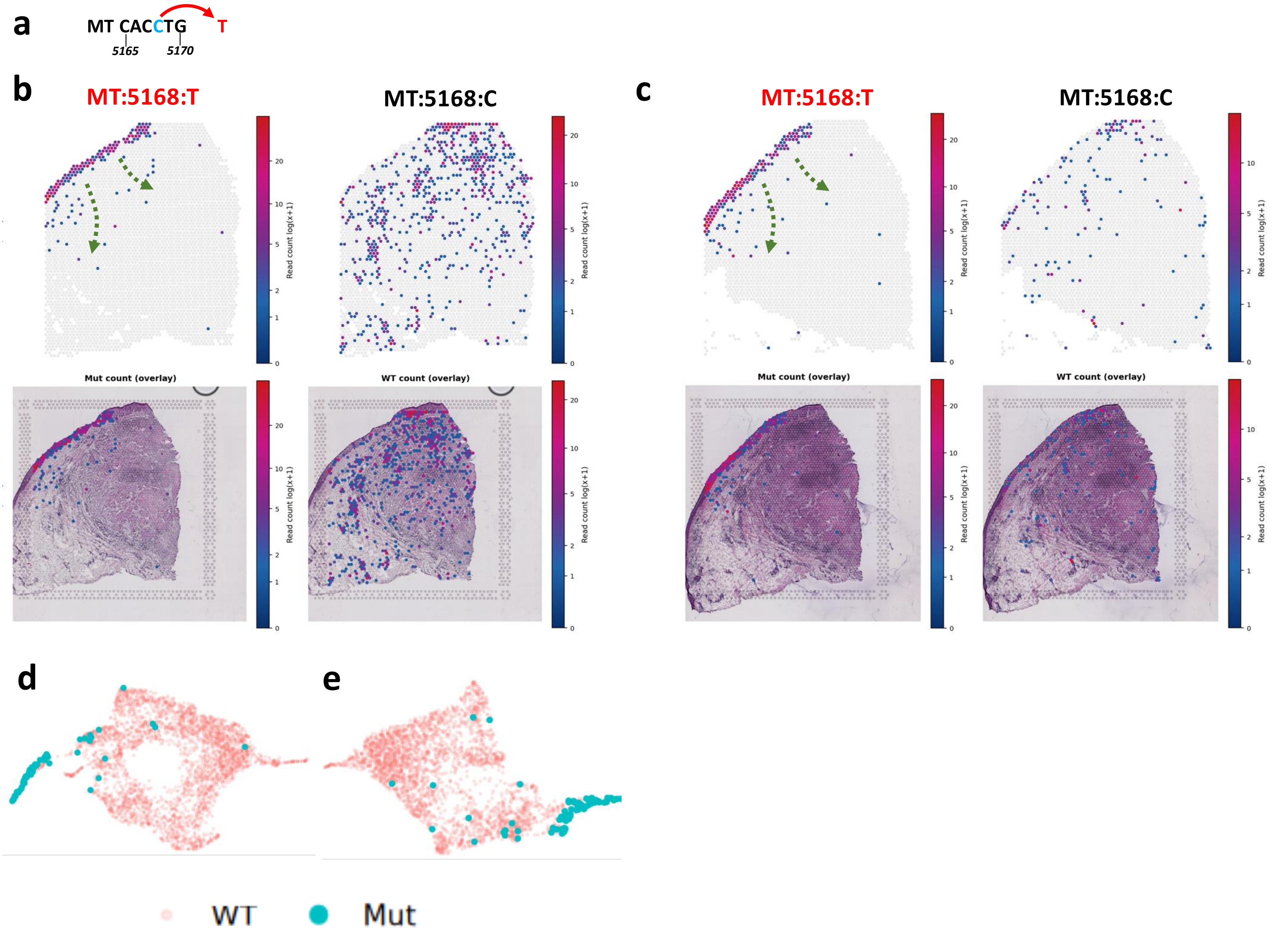
a
MT CACCTG
T
5165
5170
b
MT:5168:T
MT:5168:C
c
MT:5168:T
MT:5168:C
Read count log(x+1)
Mut count (overlay)
WT count (overlay)
d
e
WT
Mut

Figure S7

**Figure S7. Another example of previously unrecognized migration of mutation-bearing cells in skin tissue. a,** Sequence context of MT:5168:C>T, consistent with a UV-associated C>T substitution. **b-c,** Spatial distributions of the reference and mutant alleles of MT:5168:C>T in two biological replicate cSCC samples. **d-e,** UMAP of transcriptional states showing that MT:5168:C>T-positive spots are dispersed across expression space in the two biological replicate samples.
